# Heritable changes of epialleles in maize can be triggered in the absence of DNA methylation

**DOI:** 10.1101/2023.04.15.537008

**Authors:** Beibei Liu, Diya Yang, Dafang Wang, Chun Liang, Jianping Wang, Damon Lisch, Meixia Zhao

**Author notes:** Address for correspondence: Meixia Zhao.

## Abstract

*Trans*-chromosomal interactions resulting in changes in DNA methylation during hybridization have been observed in several plant species. However, very little is known about the causes or consequences of these interactions. Here, we compared DNA methylomes of F1 hybrids that are mutant for a small RNA biogenesis gene, *Mop1* (*mediator of paramutation1*) with that of their parents, wild type siblings, and backcrossed progeny in maize. Our data show that hybridization triggers global changes in both *trans*-chromosomal methylation (TCM) and *trans*-chromosomal demethylation (TCdM), most of which involved changes in CHH methylation. In more than 60% of these TCM differentially methylated regions (DMRs) in which small RNAs are available, no significant changes in the quantity of small RNAs were observed. Methylation at the CHH TCM DMRs was largely lost in the *mop1* mutant, although the effects of this mutant varied depending on the location of the CHH DMRs. Interestingly, an increase in CHH at TCM DMRs was associated with enhanced expression of a subset of highly expressed genes and suppressed expression of a small number of lowly expressed genes. Examination of the methylation levels in backcrossed plants demonstrates that TCM and TCdM can be maintained in the subsequent generation, but that TCdM is more stable than TCM. Surprisingly, although increased CHH methylation in F1 plants did require *Mop1*, initiation of the changes in the epigenetic state of TCM DMRs did not require a functional copy of this gene, suggesting that initiation of these changes is not dependent on RNA-directed DNA methylation.

## Introduction

DNA methylation is a heritable epigenetic mark involved in many important biological processes, such as genome stability, genomic imprinting, paramutation, development, and environmental stress responses [1–4]. In plants, DNA methylation commonly occurs in three cytosine contexts, the symmetric CG and CHG (where H = A, C, or T) contexts, and the asymmetric CHH context [5–7]. In Arabidopsis, *de novo* methylation at all of these three cytosine contexts is catalyzed by domains rearranged methyltransferase 2 (DRM2) through the RNA-directed DNA methylation (RdDM) pathway. In RdDM, single-stranded RNA is transcribed by RNA polymerase IV (Pol IV) and copied into double-stranded RNA by RNA-directed RNA polymerase 2 (RDR2). The dsRNA is then processed by Dicer-like 3 (DCL3) into 24-nucleotide (nt) small interfering RNAs (siRNAs), which can recruit histone modifiers and DNA methyltransferases back to the original DNA sequences to trigger methylation [3–5]. In maize, loci targeted by RdDM are primarily transposable elements (TEs) or other repeats near genes, where the chromatin is more accessible, rather than the deeply heterochromatic regions farther from genes [8, 9]. In plants, DNA methylation is maintained by different pathways depending on the location of the target sequences [6]. CG and CHG methylation are maintained during following DNA replication by methyltransferase 1 (MET1) and chromomethylase 3 (CMT3), respectively [4, 5]. CHH methylation is maintained through persistent *de novo* methylation by DRM2 through the RdDM pathway, which requires small RNAs and relatively open chromatin, or by chromomethylase 2 (CMT2) in conjunction with H3 lysine 9 dimethylation (H3K9me2) in deep heterochromatin, which does not [10].

This complex system of chromatin modification ensures that epigenetic silencing is reliably transmitted from generation to generation. However, there are situations in which that stability can be perturbed. Hybrids are an example of this because hybridization brings together two divergent genomes and epigenomes in the same nucleus. The interaction between these divergent genomes can result in both instability and transfers of epigenetic information between genomes. *Trans*-chromosomal interactions of DNA methylation between parental alleles in F1 hybrids occur in many plant species, including Arabidopsis [1, 11-14], rice [15–17], maize [18–20], pigeonpea [21], and soybean [22]. In Arabidopsis F1 hybrids, significant changes in F1 methylomes involve *trans*-chromosomal methylation (TCM) and *trans*-chromosomal demethylation (TCdM), in which the methylation level of one parental allele is altered to resemble that of the other parental allele [1, 11, 12, 21].

Small RNAs, particularly 24-nt siRNAs, are associated with the methylation changes at the regions of the genome where methylation levels differ between the two parents [1, 11, 16, 17, 23, 24]. Small RNA sequencing in Arabidopsis, maize, wheat and rice has revealed a general decrease in 24-nt siRNAs in hybrids at regions where parental siRNA abundance differs [16-18, 23, 25]. In maize, downregulation of 24-nt siRNAs following hybridization is observed in developing ears but not in seedling shoot apex [18], suggesting either the tissue type or developmental stage is important for the changes in small RNAs observed in hybrids. It has been hypothesized that siRNAs produced from the methylated parental allele can trigger *de novo* methylation of the other parental allele when the two alleles are brought together in F1 hybrids [12, 13], a process that is reminiscent of paramutation at many loci in maize [26, 27]. In Arabidopsis F1 hybrids, siRNAs from one allele are found to be sufficient to trigger methylation without triggering siRNA biogenesis from the other allele in F1 plants at TCM differentially methylated regions (DMRs) [1].

The inheritance of both TCM and TCdM in subsequent generations can be meiotically stable across many generations but varies at different loci in Arabidopsis [11, 28, 29]. In maize and soybean, parental methylation differences are inherited by recombinant inbred lines over multiple generations. However, these changes can be unstable, and are likely guided by small RNAs [22, 30]. A recent study in maize identified thousands of TCM and TCdM loci in F1 hybrids. However only about 3% of these changes were transmitted through six generations of backcrossing and three generations of selfing [31], suggesting that the methylation status of any given locus is largely determined by local sequences.

Most recent research has focused on the initiation and maintenance of overall levels of DNA methylation, but the causes and consequences of DNA methylation depend on its sequence context. In large genomes such as maize, regions distant from genes are typically maintained in a deeply heterochromatic state and cytosine methylation is primarily the CG and CHG sequence contexts. In contrast, CHH methylation, which is primarily dependent on RdDM in maize, occurs almost exclusively in regions immediately adjacent to genes, resulting in so-called “mCHH islands” [9, 32]. The result of this variation is a dramatically skewed distribution of methylated cytosines. In the maize reference genome, there are a total of 972,798,068 cytosines, out of which 18.7% and 16.4% are CG and CHG cytosines and 64.9% of which are CHH cytosines. Unlike CG and CHG cytosines, which are methylated at a high level, the level of CHH methylation is extremely low, only 2.4% genome-widely, and is largely restricted to mCHH islands. This may be due to lack of CMT2 in maize, the major chromomethylase that functions in the maintenance of CHH methylation in heterochromatin in other plants. In maize, these CHH islands are thought be the boundaries between deeply silenced heterochromatin and more active euchromatin that promote and reinforce silencing of TEs near genes [9, 32].

To address these questions, we performed high-throughput sequencing of DNA methylomes, small RNA and mRNA from F1 hybrids that were mutant for a small RNA biogenesis gene, *Mop1* (*mediator of paramutation1*), as well as their parents, wild type siblings and backcrossed progeny. *Mop1* is a sequence ortholog of *RDR2* in Arabidopsis, which is a major component of the RdDM pathway [33, 34]. In the *mop1* mutant, 24 siRNAs are dramatically reduced [8, 35], which results in a near completely removal of CHH methylation near genes [9, 36], confirming a significant role for MOP1 in *de novo* CHH methylation in maize. Our results show a global increase in CHH methylation in hybrids, but these increases are unequally distributed, leading to new and distinctive patterns of methylation. While only the low-parent (the parent with the lower methylation level) allele gained methylation in CG and CHG TCM DMRs, both the high-parent (the parent with the higher methylation level) and low-parent alleles of CHH TCM DMRs gained methylation in F1 hybrids. As has been observed in Arabidopsis, the increase in methylation in the low-parent alleles was not associated with the generation of allele-specific small RNAs at many genomic loci, suggesting that small RNAs from one allele are sufficient to trigger methylation in the other allele, but are not always sufficient to trigger Pol IV transcription of the target allele. Interestingly, these CHH TCM DMRs were associated with the enhanced expression of a subset of highly expressed genes and suppressed expression of a subset of lowly expressed genes.

Changes in CG and CHG methylation were often retained in the backcrossed generation, a process that did not require MOP1. Heritable changes in CHH methylation were more complex. The increase in CHH methylation in both the highly methylated and lowly methylated alleles was lost in backcrossed 1 (BC1) plants, even at loci where both alleles were present, suggesting that the global increase we observed in the F1 is a function of heterosis, rather than an interaction between each pair of heterozygous epialleles. However, new methylation added to the low methylation allele could be transmitted to the BC1 plants, even in progeny of plants that were *mop1* mutant and that lacked MOP1-dependent siRNAs. This suggests that the transfer of the epigenetic state from high CHH alleles to low CHH alleles, as well as the maintenance of this altered state in the gametophyte does not require MOP1.

## Results

### CHH methylation level is increased globally in hybrids

To understand the initiation of DNA methylation, we crossed *mop1* heterozygous plants in the Mo17 and B73 backgrounds to each other (Mo17;*mop1-1/+* × B73;*mop1-1/+*) to generate F1 hybrid *mop1* mutants (Mo17/B73;*mop1-1/mop1-1*, designated as *mop1*F1) and their hybrid homozygous wild type siblings (Mo17/B73;*+/+*, designated as WTF1) (Fig 1A). We next performed whole genome bisulfite sequencing (WGBS) of the two parental genotypes (Mo17 and B73) and the two F1 hybrids (WTF1 and *mop1*F1) (S1 Table). The overall methylation levels of B73 (25.1%) and Mo17 (25%) were similar. We observed a substantial increase in overall methylation levels in WTF1 hybrids (30%) compared to the two parents (25%), as has been noted previously in both Arabidopsis and maize (S1 Fig and S2 Table) [1, 31]. The increased methylation was primarily driven by the increased CHH methylation, while CG and CHG were not dramatically changed (Fig 1B, S1 Fig and S2 Fig). In both parents and WTF1, the overall levels of CHH methylation tend to be higher in chromosomal arms, likely because there are more mCHH islands near genes in the ends of chromosomes [9]. Interestingly, although the *mop1* mutation reduces CHH methylation [36], the overall level of CHH methylation in *mop1*F1 was still higher than the two wild type parents (Fig 1B and S2 Fig), suggesting that a significant portion of the increased *de novo* CHH methylation in F1 hybrid plants does not require classical RdDM.

**Fig 1.**
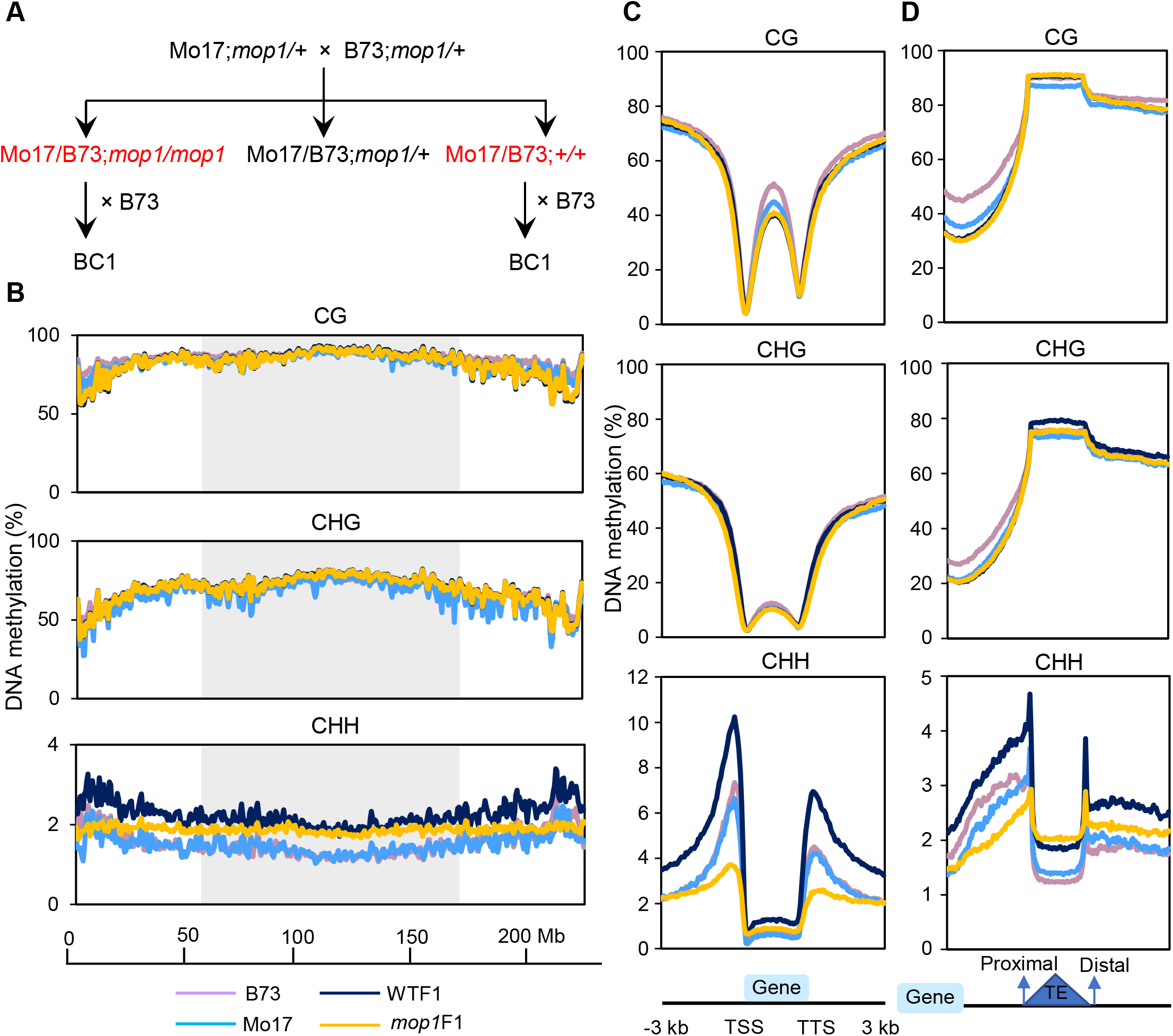
CHH methylation level is globally increased in hybrids. (**A**) Genetic strategy to construct wild type F1 (WTF1), *mop1* mutant F1 (*mop1*F1), and backcross1 (BC1). **(B)** The distribution of CG, CHG, and CHH methylation on chromosome 5. Methylation levels were measured in 1 Mb windows with 500 kb shift. The shaded boxes represent pericentromeric regions. **(C)** Patterns of methylation in and flanking genes. **(D)** Patterns of methylation in and flanking TEs. DNA methylation levels were calculated in 50 bp windows in the 3 kb upstream and downstream regions of the genes/transposable elements (TEs). Each gene/TE sequence was divided into 40 equally sized bins to measure the gene/TE body methylation. Bin sizes differ from gene/TE to gene/TE because of the different lengths of genes/TEs. The methylation levels of TEs were orientated into proximal and distal ends depending on the flanking genes of TEs. Methylation for each sample was calculated as the proportion of methylated C over total C in each sequence context (CG, CHG, and CHH, where H = A, T, or C) averaged for each window. The average methylation levels were determined by combining two biological replicates for each genotype.

Previous research had shown that *mop1* mutants primarily affect mCHH islands near active genes [8, 9]. Therefore, we plotted DNA methylation levels of CG, CHG and CHH within gene bodies, 3 kb upstream of TSSs (transcription start sites) and 3 kb downstream of TTSs (transcription termination sites). In genes, we observed similar patterns with respect to the methylation levels of CG and CHG between parents and F1 hybrids. In contrast, the methylation levels of CHH cytosines both upstream and downstream of genes were dramatically increased in WTF1 plants, and dramatically reduced in the *mop1*F1 mutants relative to the two parents (Fig 1C). We next determined CG, CHG and CHH methylation levels within TE bodies and their flanking regions. The region flanking the distal edge of TEs relative to genes generally had higher levels of CG and CHG methylation than did the region flanking their proximal edge. CHH methylation was increased in WTF1 hybrids across TE bodies and flanking regions relative to the parents, particularly at the two edges of TEs. In line with previous observations [9], CHH methylation level at the proximal edge and the adjacent flanking regions of TEs in *mop1*F1 was lower than that in the two parents. In contrast, the CHH methylation level at the distal edge of TEs and the adjacent flanking regions in *mop1*F1 was only marginally reduced relative to WTF1, and was still higher than that in the parents (Fig 1D). In the body of TEs, the increase in CHH methylation triggered by hybridization was unchanged, or even increased in *mop1* mutants.

Together, these data suggest that MOP1 is particularly important for CHH methylation of the ends of TEs that are near genes, along with the region between the TE and the gene. Outside of those regions, it appears that MOP1 is not required for a significant portion of the increased CHH methylation in F1 plants. The net effect is a strong effect of *mop1* on CHH islands, but a much reduced effect on overall changes in DNA methylation seen in the F1 generation.

### Levels of CHH methylation of both high– and low-parent (parents with the higher and lower methylation levels) alleles are increased at TCM DMRs in the F1 hybrids

We identified DMRs between the two parents, Mo17 and B73, in our data set. Here we referred to these DMRs as parental DMRs, which can be Mo17 or B73 hyper DMRs, indicating that either Mo17 or B73 has a significantly higher level of DNA methylation (Fig 2A). In total, we identified 7,107 CG, 9,045 CHG, and 13,307 CHH DMRs between the two parents (Fig 2B and S3 Table). CHH DMRs were typically shorter than CG and CHG DMRs (S3 Fig and S3 Table). The B73 genome had more CG and CHG hyper DMRs, and the Mo17 genome had more CHH hyper DMRs, which is consistent with the observation that B73 had higher overall CG and CHG methylation and Mo17 had higher overall CHH methylation at these DMRs (Fig 2B and 2C), as has been noted previously [37]. We also found that CG and CHG DMRs were more overlapped with each other than each one was with CHH DMRs (Fig 2D), consistent with previous observations that CHH methylation is often found in mCHH islands immediately up and downstream of genes [9, 32]. Out of the 13,307 CHH DMRs, 52% were located within or near genes, particularly 2 kb upstream and downstream of genes (43%), which was significantly higher than the values for CG (27%) and CHG (18%) in these regions (*P* < 0.0001, χ^2^ test) (Fig 2E). Given that TEs are the primary targets of DNA methylation and maize genes are frequently adjacent to TEs [3, 9], we compared the different classes of TEs overlapping DMRs within the 2 kb flanking regions of genes. Not surprisingly given their distribution within genomes, we found that terminal inverted repeat (TIR) DNA transposons were more enriched in CHH DMRs than they were in CG and CHG DMRs within 2 kb of genes (Fig 2E and 2F).

**Fig 2.**
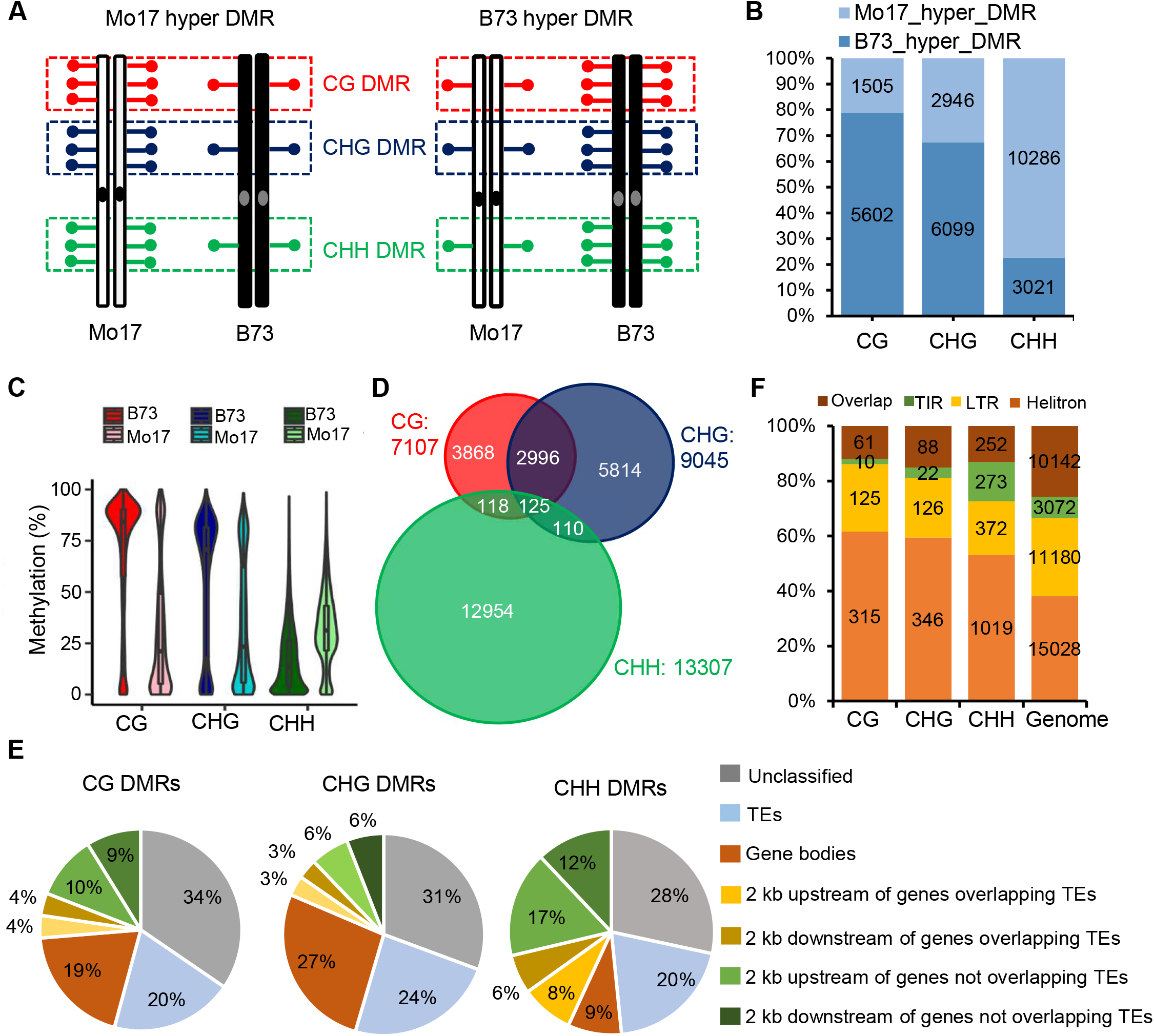
Parental CHH DMRs are largely located within 2 kb flanking regions of genes. (**A**) Definition of B73 hyper DMRs (higher methylation in B73) and Mo17 hyper DMRs (higher methylation in Mo17) between parents. Red, green, and blue dots represent CG, CHG, and CHH methylation, respectively. **(B)** B73 has more CG and CHG hyper DMRs, and Mo17 has more hyper CHH DMRs. **(C)** B73 has higher methylation levels at CG and CHG DMRs, and Mo17 has higher methylation levels at CHH DMRs. **(D)** CG and CHG DMRs were more overlapped with each other than each one was with CHH DMRs. **(E)** The distribution of CG, CHG and CHH parental DMRs. 2 kb up and downstream of genes overlapping TEs indicate the DMRs overlap TEs within the 2 kb flanking regions of genes. **(F)** The types of TEs in the categories of 2 kb up and downstream of genes overlapping TEs in (E). DMRs, differentially methylated regions.

Next, we examined the methylation levels of these parental DMRs in the F1 hybrids. Following previously published studies, we compared the methylation levels of WTF1 to the mid-parent value (MPV, the average of the two parents) and classified changes as being a consequence of TCM (*trans*-chromosomal methylation), TCdM (*trans*-chromosomal demethylation), or NC (no change) (Fig 3A) [1]. A majority of parental DMRs (∼75%) did not significantly change their methylation levels in the WTF1 hybrids, and most of these unchanged DMRs were in TEs and unclassified regions (Fig 3A and S4 Fig). However, when single nucleotide polymorphisms (SNPs) were used to distinguish methylation in each of the two parental genomes, many of these NC DMRs (CG 53.8%, CHG 52.9%, and CHH 51.4%) in WTF1 were revealed to have lost methylation at the high-parent allele and gained methylation at the low-parent allele, which resulted in no significant changes in overall methylation levels between the hybrids and parents, suggesting that methylation interaction still occurs in these NC DMRs (S5 Fig). Of the remaining 25% parental DMRs that were significantly changed in F1 hybrids, 18.7% were TCM, and 6.8% were TCdM (Fig 3A). We then compared allele specific methylation levels of these regions between B73 and Mo17. Given that these two inbred genomes are highly polymorphic, we were able to compare allele specific methylation at 2,459 (57%) of the TCM and 915 (59%) of the TCdM DMRs. At TCM DMRs, WTF1 had higher methylation levels at all three cytosine contexts (Fig 3B). The increased methylation at CG and CHG in these wild type F1 plants was primarily due to the increased methylation in the parental allele that had the lower level of methylation. In contrast, CHH methylation levels of both the high– and low-parent alleles were substantially increased in WTF1 at these TCM DMRs (Fig 3B). At TCdM DMRs, the reduction of methylation was primarily due to the decreased methylation of the high-parent allele in all of the three cytosine contexts (Fig 3C).

**Fig 3.**
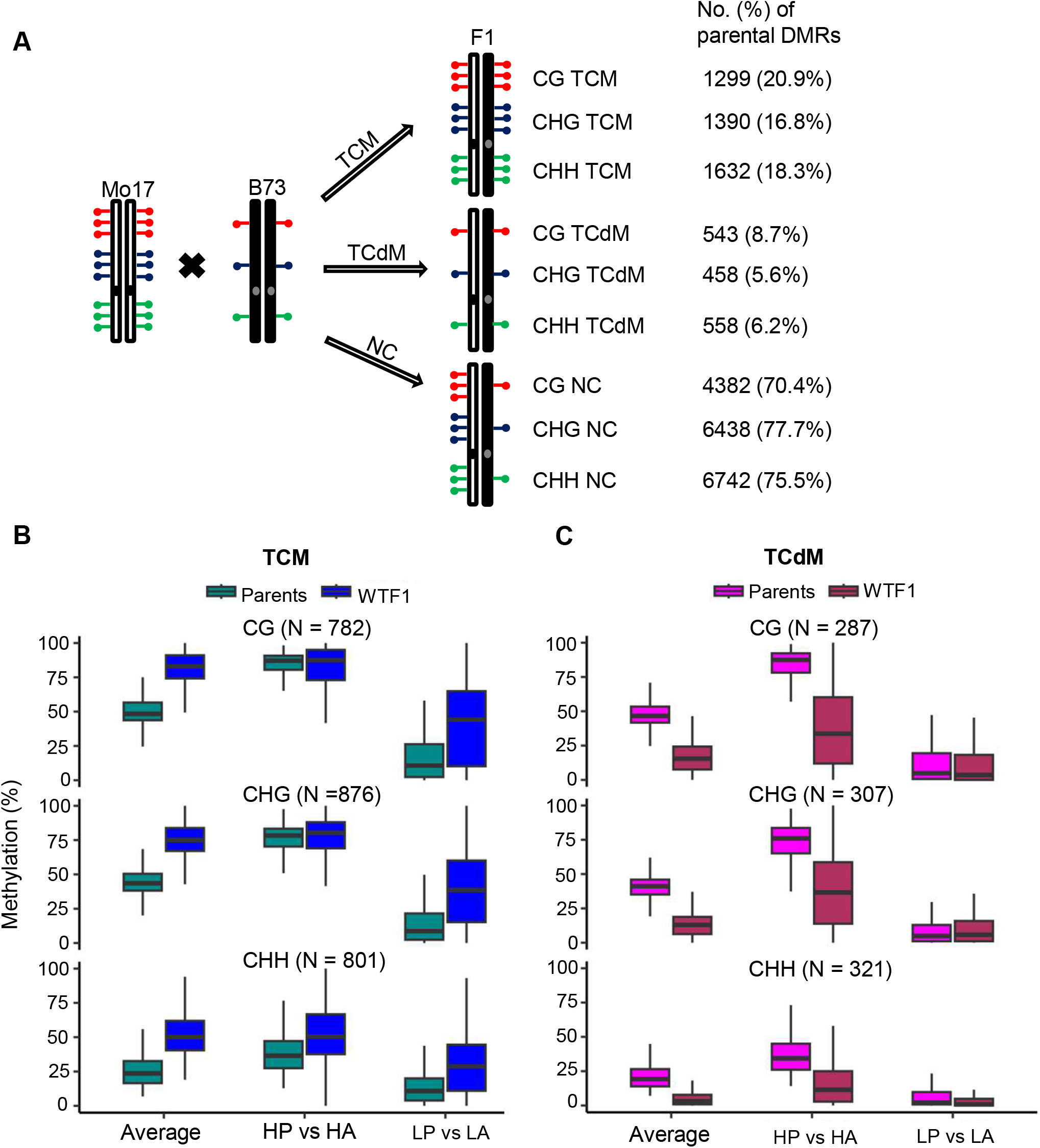
The levels of CHH methylation of both high– and low-parent alleles are increased at TCM DMRs in the F1 hybrids. (**A**) Identification of TCM, TCdM, and unchanged (NC) DMRs between WTF1 and parents. **(B)** Comparisons of CG, CHG and CHH methylation at TCM DMRs in parents and WTF1. **(C)** Comparisons of CG, CHG and CHH methylation at TCdM DMRs in parents and WTF1. HP, high parent (parent with higher methylation). HA, high-parent allele in F1. LP, low parent (parent with lower methylation). LA, low-parent allele in F1. Average means the average between the two parents, or between the two alleles in WTF1 and *mop1*F1. DMRs, differentially methylated regions. TCM, *trans*-chromosomal methylation. TCdM, *trans*-chromosomal demethylation.

### Methylation of CHH TCM DMRs is dramatically reduced in the *mop1* mutant

To shed light on the effects of the loss of *Mop1*-dependent small RNAs at TCM and TCdM DMRs, we examined their methylation levels in *mop1*F1 mutant plants. Only 99 (8.6%) of 1,147 CG and 144 (11.2%) of 1,284 CHG TCM DMRs significantly changed their methylation levels in *mop1*F1 mutants. In contrast, methylation levels of 90.7% (1,031 out of 1,137) CHH TCM DMRs were significantly changed in *mop1*F1 (Fig 4A). Consistent with our global analysis, the CHH DMRs that were significantly changed in *mop1* were primarily located in the 2 kb flanking regions of genes (S6 Fig). As expected, methylation of all the three sequence contexts at these TCM DMRs were largely reduced in *mop1*F1 mutants, particularly in the CHH context, in which the methylation level in *mop1*F1 plants was even lower than the low parent (Fig 4B). This suggests that in these regions, but not the genome as a whole, the additional methylation in F1 wild type plants is lost altogether. Not surprising, given that the methylation of TCdM DMRs was already very low, we did not observe significant changes in methylation at TCdM DMRs in the *mop1*F1 mutants (Fig 4C).

**Fig 4.**
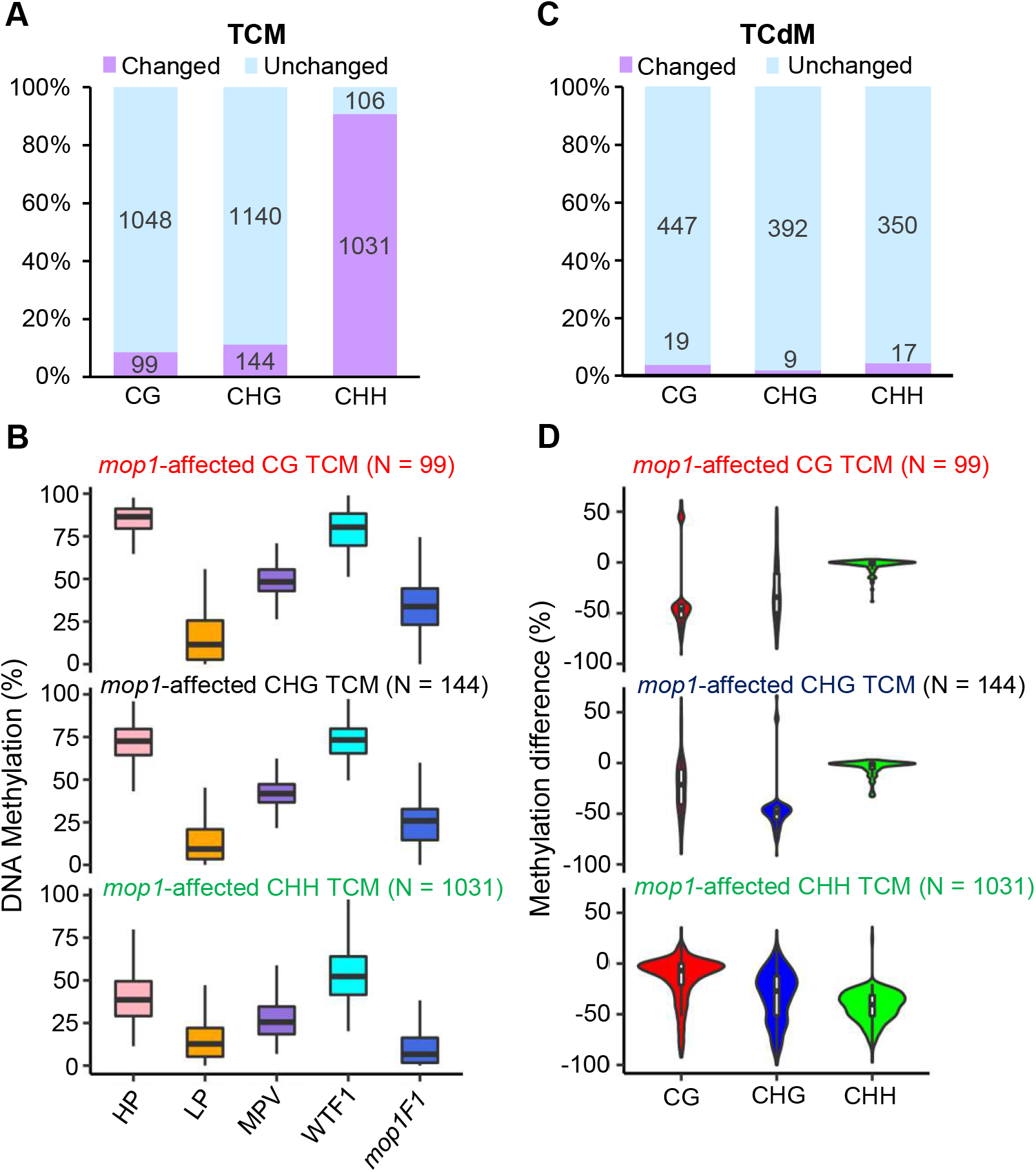
The *mop*1 mutation primarily removes the methylation of CHH TCM DMRs. (**A**) Number of CG, CHG and CHH TCM DMRs affected by the *mop1* mutation. **(B)** Comparison of methylation levels at the *mop1*-affected CG, CHG, and CHH TCM DMRs. **(C)** Number of CG, CHG and CHH TCdM DMRs affected by the *mop1* mutation. **(D)** Examination of the methylation changes in the other two cytosine contexts at the *mop1*-affected CG, CHG, and CHH TCM DMRs. The top panel shows the methylation changes in CHG and CHH sequence contexts for the 99 *mop1*-affected CG TCM DMRs. The middle panel shows the methylation changes in CG and CHH sequence contexts for the 144 *mop1*-affected CHG TCM DMRs. The bottom panel shows the methylation changes in CG and CHG sequence contexts for the 1031 *mop1*-affected CHH TCM DMRs. HP, high parent (parent with higher methylation). LP, low parent (parent with lower methylation). MPV, the middle parent value. DMRs, differentially methylated regions. TCM, *trans*-chromosomal methylation. TCdM, *trans*-chromosomal demethylation.

Previous research has demonstrated that loss of methylation in mCHH islands results in additional loss of CG and CHG methylation [9]. We found that out of the 118 CG DMRs that were significantly changed in *mop1*F1 mutants relative to their wild type siblings, 37 (31.4%) were also CHG DMRs, but only 3 (2.5%) were CHH DMRs. Similarly, only 32 (20.9%) and 9 (5.9%) of the CHG DMRs that were changed in *mop1* were CG and CHH DMRs, respectively. Out of the 1,048 *mop1*-affected CHH DMRs, 72 (6.7%) and 181 (17.3%) were also CG and CHG DMRs (S4 Table). A similar pattern was observed for the *mop1*-affected CHG DMRs, in which we detected changes in CG but no changes in CHH methylation. For *mop1*-affected CHH DMRs, we saw no change in CG but a substantial change in CHG (Fig 4D). Together these data suggest that *mop1* mutation primarily prevents the methylation of CHH TCM DMRs, and that a loss of CHH methylation in *mop1* can result in additional loss of CHG, but not CG methylation.

### Small RNAs from one allele are sufficient to trigger methylation of the other allele at a majority of CHH TCM DMRs in F1 hybrids

Because small RNAs are the trigger for *de novo* DNA methylation [4, 5], we next asked whether the difference in methylation during hybridization is caused by differences in small RNAs. We proposed two hypotheses with respect to siRNAs at the CHH TCM DMRs. As shown in Fig 5A, in the first hypothesis, small RNAs are produced from one allele and trigger increases in methylation at the high-parent allele and *de novo* methylation in low-parent allele without triggering production of new, allele specific small RNAs from that allele. In the alternative hypothesis, once methylation is triggered in the low-parent allele, it becomes competent to produce its own, allele specific small RNAs, which may in turn act to enhance at the high-parent allele. To distinguish between these hypotheses, we performed small RNA sequencing from the same plants that were used for DNA methylation analysis (S1 Table). Because of the increase in the apparent number of 22-nt siRNAs in *mop1* mutants caused by normalization following the loss of most 24-nt small RNAs in *mop1* mutants, the small RNA values were adjusted to total abundance of all mature microRNAs following previously described protocols [35]. As was expected, 24-nt siRNAs were the most abundant siRNAs in all the sequenced wild type samples. Overall, despite the dramatic increase we observed in CHH methylation in the hybrids (Fig 1B), no significant differences in small RNAs were observed between the WTF1 hybrids and parents (Fig 5B). The *mop1* mutation substantially reduced 24-nt siRNAs, particularly in the mCHH island regions near TSSs and TTSs (Fig 5B and S7 Fig). Next, we compared 24-nt siRNAs generated from the high parent and low parent. We detected 24-nt uniquely mapped siRNAs in 795 CG (11.2% of the total), 700 CHG (7.7%), and 5,070 CHH (38.1%) parental DMRs. Consistent with their role in methylation, on average, the high parent harbored significantly more 24-nt siRNAs than the low parent (Fig S8). This is also true for TCM, TCdM, and NC DMRs when analyzed separately (S9 Fig).

**Fig 5.**
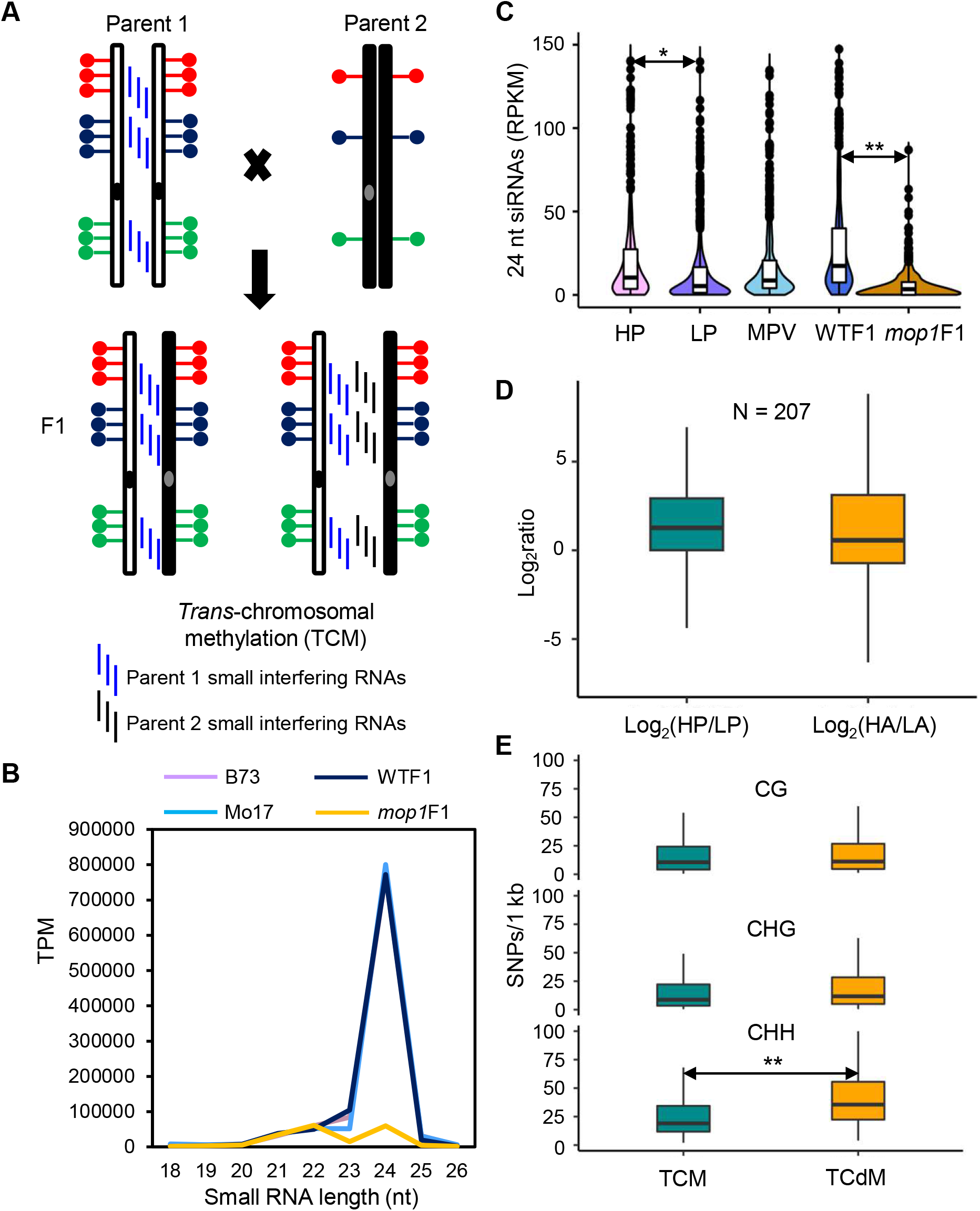
Small RNAs produced from one parent are sufficient to trigger new methylation of the other allele in hybrids. (**A**) Two hypothetical models of small RNA biogenesis in F1 at CHH TCM. **(B)** Expression values of small RNAs in parents, WTF1 and *mop1*F1. TPM, transcripts per million uniquely mapped reads. The small RNA values were adjusted to total abundance of all mature microRNAs following the previous research [35]. **(C)** The abundance of 24-nt siRNAs at the *mop1-*affected CHH TCM DMRs. HP, high parent (parent with higher methylation). LP, low parent (parent with lower methylation). MPV, the middle parent value. RPKM, 24-nt siRNA reads per kilobase (DMR length) per million uniquely mapped reads. **(D)** Ratios of 24-nt siRNAs of the high parent to the low parent, and of the high-parent allele to the low-parent allele at the *mop1*-affected CHH TCM DMRs. **(E)** Number of single nucleotides polymorphisms between TCM and TCdM. **, *P* < 0.01, *, *P* < 0.05, Student’s *t* test. DMRs, differentially methylated regions. TCM, *trans*-chromosomal methylation. TCdM, *trans*-chromosomal demethylation.

To test whether the increase in methylation in WTF1 plants was due to an increase in 24-nt small RNAs, we compared the abundance of 24-nt siRNAs between WTF1 and the MPV. Although 24-nt siRNAs were increased at CHH TCM DMRs in WTF1 hybrids, this increase was not significant (Fig 5C and S9 Fig). We then analyzed allele-specific expression of siRNAs in F1 hybrids. Because only uniquely mapped reads with SNPs can be used to access the allele specific expression of siRNAs and because the length (24-nt) of siRNAs is very short, we were able to obtain data from only 207 CHH TCM DMRs that had enough information to compare allele-specific expression. There was no significant difference between the ratio of 24-nt siRNAs of the high-parent allele to the low-parent allele in F1 hybrids and that of the high parent to the low parent in the parents (Fig 5D). Among these 207 CHH TCM DMRs, 53 had siRNAs expressed from only the high parent. Of these, 34 (64.2%) had siRNAs still produced from the high-parent allele in WTF1. Out of the remaining 154 CHH TCM DMRs, 104 expressed more siRNAs from the high parent, out of which, 65 (62.5%) still had more siRNAs expressed from the high-parent allele than the low-parent allele in WTF1. These data suggest that the increased methylation at CHH TCM DMRs is not caused by an increase in siRNAs from the newly methylated allele, which favors the hypothesis that small RNAs produced from one allele trigger methylation of the other allele in *trans,* but that the newly methylated allele is not itself a source of small RNAs.

RdDM triggered by small RNAs depends on the similarity of the small RNAs and their targets. Thus, the sequence variation between the two alleles may affect small RNA targeting and ultimately, methylation. To test this, we compared the SNPs between TCM and TCdM. As shown in Fig 5E, no significant differences in SNP enrichment were observed when comparing TCM and TCdM at CG and CHG DMRs. In contrast, CHH TCdM DMRs had significantly more SNPs than did CHH TCM DMRs, suggesting that more genetic variation at CHH TCdM DMRs hinders targeting of one allele by small RNAs from the other allele.

### CHH methylation of sequences flanking genes can be associated with either suppressed or enhanced expression of neighboring genes

Given the variation in DNA methylation we observed in the parental lines and F1 hybrids (Fig 2), we compared the expression values of 51 genes involved in the RdDM pathway among these genotypes. We detected eight RdDM genes differentially expressed between B73 and Mo17, all of which showed significantly higher expression in the Mo17 genome (S10 Fig and S5 Table), which may contribute to the greater abundance of CHH methylation in the Mo17 genome (Fig 2B and 2C). In addition, we identified six RdDM pathway genes differentially expressed between the F1 hybrids and the MPV, and all of them had higher expression in the F1 hybrids (S10 Fig and S6 Table), suggesting that the RdDM pathway is more active in hybrids.

DNA methylation is generally associated with repression of transcription, particularly when the methylation is in the promoter regions of genes [38–40]. However, previous analysis of the maize methylome suggests that the reverse is true of CHH islands. One interpretation of this observation is that because CHH methylation is an active process that requires relatively open chromatin, increased gene expression may permit more efficient RdDM, resulting in higher levels of methylation [9, 32]. If this were the case, one would expect that allele specific increases in expression in F1 plants would result in increased CHH methylation of TEs near those genes. Alternatively, it is possible that additional CHH methylation could under some circumstance, result in decreased expression in F1 plants. To understand the relationship between CHH methylation and gene expression, we investigated the correlation between a subset of CHH TCM DMRs with expression of genes that flank them. As shown in Fig 6A, for the Mo17 CHH TCM DMRs, whose methylation is transferred from Mo17 to B73, if methylation suppresses gene expression, because the Mo17 parent has higher methylation, we expect the Mo17 allele to have a lower level of expression. After hybridization, if the B73 allele gains methylation, it would be expected to produce less transcript. If this is the case, we would expect to see the ratio of gene expression of B73 to Mo17 in the F1 hybrids to decrease relative to the ratio of expression of these alleles in the parents (Fig 6A, left panel). In contrast, if CHH methylation promotes gene expression or if it responds passively to increased expression, we would predict an increase in the ratio of gene expression of B73 to Mo17 in F1 hybrids associated with a relative increase in expression of B73 (Fig 6A, right panel).

**Fig 6.**
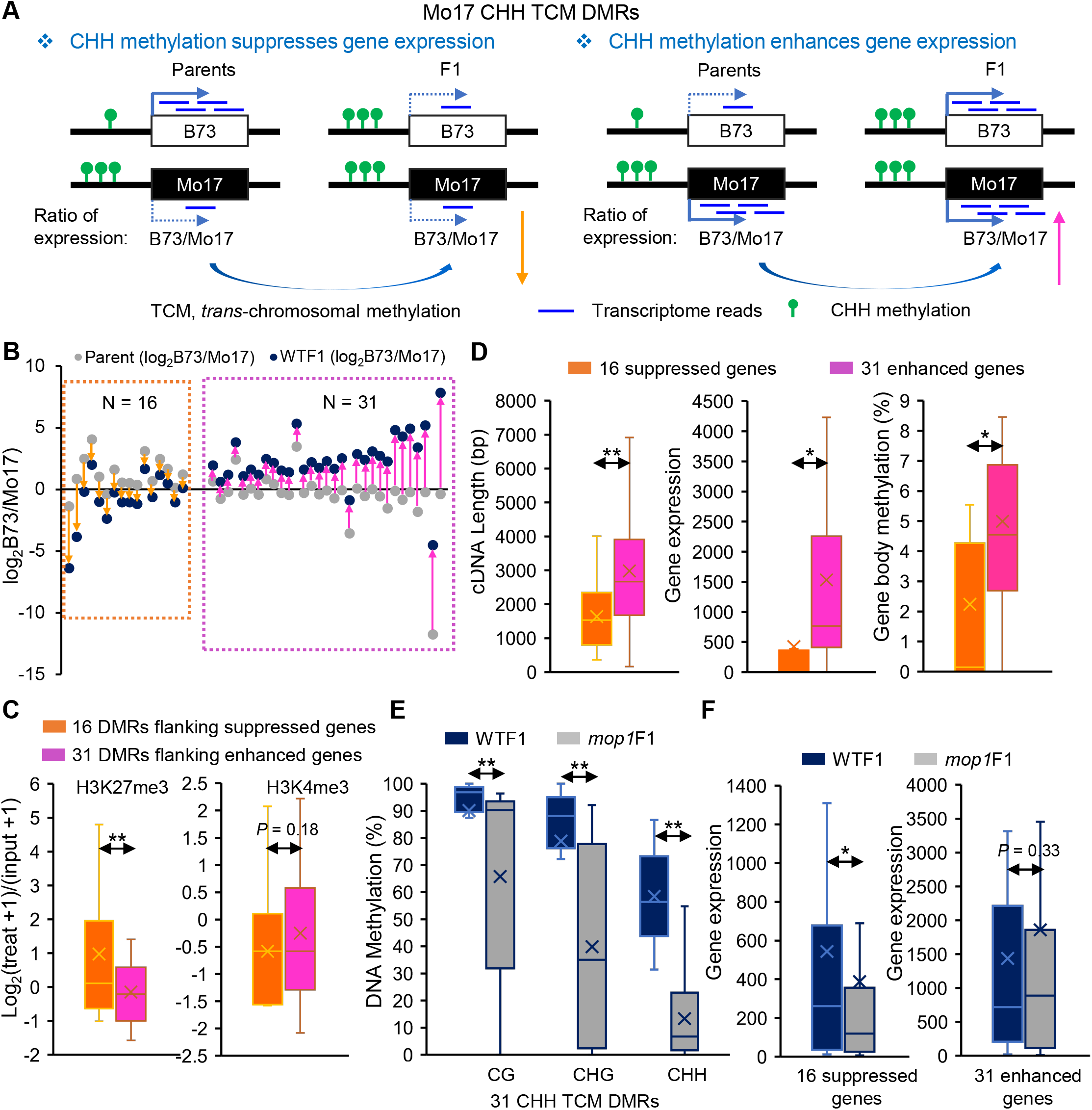
CHH methylation is associated with both suppressed and enhanced expression of their flanking genes. (**A**) Two possible scenarios of the effects of CHH methylation on gene expression. Only examples of Mo17 CHH TCM DMRs are shown. **(B)** Number of genes identified based on the models in (A). The ratios of expression values of B73 to Mo17 between the parents and WTF1 were used to distinguish the scenarios. **(C)** Histone modification of 16 CHH TCM DMRs that are associated with the suppressed expression of flanking genes and 31 CHH TCM DMRs that are associated with the enhanced expression of flanking genes. **, *P* < 0.01, Student’s *t* test. **(D)** Gene properties of the 16 suppressed and 31 enhanced genes. **, *P* < 0.01, *, *P* < 0.05, Student’s *t* test. **(E)** Methylation changes in *mop1* mutants at the 31 CHH TCM DMRs that are associated with the enhanced expression of flanking genes. **, *P* < 0.01, Student’s paired *t* test. **(F)** Gene expression changes of the 16 suppressed and 31 enhanced genes in *mop1* mutants. *, *P* < 0.01, Student’s paired *t* test. DMRs, differentially methylated regions. TCM, *trans*-chromosomal methylation.

We focused on 442 Mo17 hyper DMRs with available data on allele-specific methylation in F1, 172 of which also had available allele-specific expression data, and then looked for genes whose allele-specific expression changed significantly in F1 relative to the parents. Of the genes flanking the 172 Mo17 hyper DMRs, 126 (73%) associated with those DMRs showed no significant change in relative expression. For 16 genes, the ratio of B73 to Mo17 expression was decreased, and for 31 genes, the ratio was increased in the F1 hybrids (Fig 6B), suggesting that CHH methylation can be associated with both suppressed and enhanced gene expression. Next, we asked whether variation in expression of genes is associated with variation in histone modifications. The 16 DMRs that were associated with suppressed gene expression were significantly more enriched for H3K27me3 and more depleted of H3K4me3 than the 31 DMRs that were associated with enhanced gene expression in B73 (Fig 6C and S11C Fig) [41]. The 31 genes that seemed to be enhanced by CHH methylation were typically longer, more highly expressed, and with higher gene body methylation than the 16 suppressed genes in Mo17 (Fig 6D and S11D Fig).

Because the *mop1* mutation results in reduced methylation in mCHH islands near genes [9, 36], we wanted to determine whether removal of methylation of the 16 and 31 DMRs in the *mop1* mutant changed the expression of their flanking genes. We compared the methylation levels of CHH, CG, and CHG in the 16 and 31 CHH TCM DMRs. Because CG and CHG methylation at these 16 suppression-associated CHH DMRs were not available in the *mop1* mutant, we were only able to examine methylation of the 31 enhanced-associated CHH DMRs in the *mop1* mutant. As expected, in the *mop1* mutant, CHH methylation was greatly reduced in these 31 DMRs, as was CG and CHG methylation (Fig 6E), which echoes previous research [9]. However, this reduction in CHH methylation did not have a significant effect on the expression of the 31 genes that seemed to be promoted by CHH methylation (Fig 6F). These data suggest that variations in CHH methylation are a consequence, rather than a cause of variation in gene expression.

### Most newly induced CG and CHG DMRs lose methylation, and most newly induced CHH DMRs gain methylation in F1 hybrids

In addition to examining changes in methylation of the parental DMRs, we also investigated the newly induced DMRs in F1 hybrids that were not differentially methylated in the parents. These newly induced DMRs can either gain or lose methylation at an allele relative to both parents (S12A and S12B Fig). A total of 715 CG, 1,149 CHG, and 3,876 CHH new DMRs were identified (S13 Fig). These newly induced DMRs were equally distributed as hyper or hypo DMRs relative to both the B73 and Mo17 genomes (S13C and S13D Fig), which is different from the parental DMRs, which were enriched for CHH methylation in the Mo17 genome (Fig 2B). The newly induced DMRs at CG and CHG sequence contexts largely overlapped with TEs (S13E and S13F Fig), confirming that TEs are the most frequent targets of DNA methylation. Next, we compared the allele specific methylation of these newly induced DMRs. Because the two parents were methylated at the similar levels to those at these newly induced DMRs, we defined the high or low parent as the parental allele that was changed in the F1, so the low parent would be the allele whose methylation was reduced in the F1. We found that the majority of newly induced CG (89%, 558 out of 627) and CHG (75%, 918 out of 1,231) DMRs followed the model in S12B Fig, in which one parental allele loses methylation in F1 hybrids (S12D Fig). Interestingly, the majority of newly induced CHH (92%, 2,959 out of 3,230) DMRs followed the model in S12A Fig, in which one parental allele gains methylation in F1 hybrids (S12C Fig), suggesting a distinction between CHH and CG and CHG methylation. We also compared the small RNAs at these newly induced CHH DMRs and did not observe any significant changes in small RNAs between the two parents that had similar methylation levels, or between the hybrids and parents (S14 Fig).

### Initiation of the changes in the epigenetic state of targets of *trans*-chromosomal CHH methylation does not require *Mop1*

We next wanted to determine whether the methylation or demethylation triggered in F1 can be maintained in subsequent generations. To test this, we backcrossed WTF1 (Mo17/B73;+/+) and *mop1*F1 (Mo17/B73;*mop1*/*mop1*) with B73 and obtained backcrossed (BC1) plants for WGBS (Fig 1A). We first analyzed the overall methylation differences between WTF1 and WTBC1. To determine whether changes had been heritably transmitted, we set a cut off for a lack of a change from WTF1 to WTBC1 as <10% change in methylation for CG and CHG and <5% for CHH methylation. We found that approximately 25% of CG and 26% of CHG TCM DMRs met this cut off, as did 11% CHH TCM DMRs. Interestingly, the CG (35%), CHG (44%) and CHH (38%) TCdM DMRs all had higher percentages of DMRs that met the threshold of differences between F1 and BC1 (S7 and S8 Table), suggesting that TCdM DMRs are more heritable.

To better understand the inheritance of newly acquired DNA methylation, we focused specifically on the inheritance of TCM DMRs. Because all of the sequenced BC1 plants were backcrossed individuals derived from the cross of F1 with B73, we separately analyzed B73 and Mo17 TCM DMRs. For B73 TCM DMRs (Fig 7A and Fig 8A), in which the Mo17 allele has acquired new methylation in F1, BC1 plants could be either homozygous or heterozygous for B73. Among the homozygous BC1 plants, all should have the native level of B73 methylation because it was the Mo17 allele whose methylation was changed in the F1 at these DMRs. Similarly, BC1 plants that were heterozygous for B73 and the newly converted Mo17 allele would be expected to remain hypermethylated because Mo17 was still in the presence of the B73 allele in these plants (Fig 7A and Fig 8A). For CG and CHG methylation, this is what we observed. Methylation at these DMRs in all BC1 progeny, both homozygotes and heterozygotes were at similar levels as the F1 heterozygotes (Fig 7C, 7D, S15C and S15D Fig). Whether or not the F1 plant was *mop1* or not did not affect the heritability of the added methylation in these cases.

**Fig 7.**
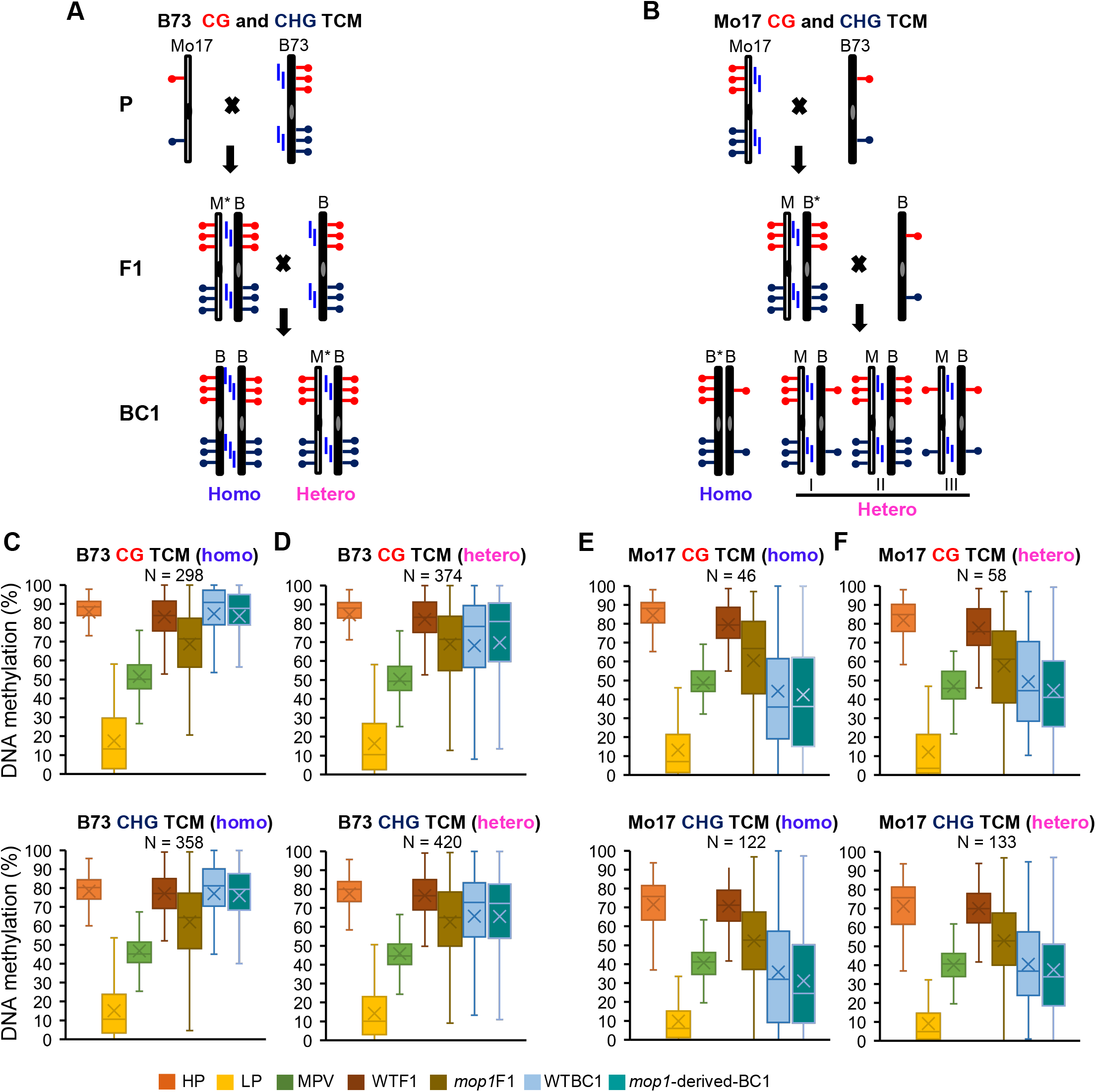
Newly triggered methylation at CG and CHG TCM DMRs in F1 plants is maintained in the next generation. (**A**) Hypothetical model of maintenance of B73 CG and CHG TCM DMRs. Asterisks denote the newly converted (methylated) allele. **(B)** Hypothetical model of maintenance of Mo17 CG and CHG TCM DMRs. **(C)** Methylation changes of homozygous B73 CG and CHG TCM DMRs. **(D)** Methylation changes of heterozygous B73 CG and CHG TCM DMRs. **(E)** Methylation changes of homozygous Mo17 CG and CHG TCM DMRs. **(F)** Methylation changes of heterozygous Mo17 CG and CHG TCM DMRs. DMRs, differentially methylated regions. TCM, *trans*-chromosomal methylation. Homo, homozygous. Hetero, heterozygous. WTBC1, Mo17/B73;+/+ × B73. *mop1*-derived BC1, Mo17/B73;*mop1/mop1* × B73.

Inheritance of CHH methylation was complicated by the fact that in each case, both alleles in the F1 had elevated methylation relative to parents. Thus, it was possible that each allele or both would return to their original level of methylation following the backcross. With respect to CHH methylation, the BC1 B73 homozygotes showed levels of methylation similar to the B73 (HP) parent, rather than the F1 (Fig 8C and S16C Fig), suggesting that the enhanced methylation in the F1 that resulted from the interaction between B73 and Mo17 had been reduced in the BC1. In the heterozygous BC1s, the overall level of methylation was more similar to the MPV than to the heterozygous F1s (Fig 8D and S16D Fig). This observation suggests that the elevated level of methylation is a consequence of the hybrid genomes, and not just an interaction between alleles, and that the elevated levels of methylation at Mo17 in the F1 was not heritable.

**Fig 8.**
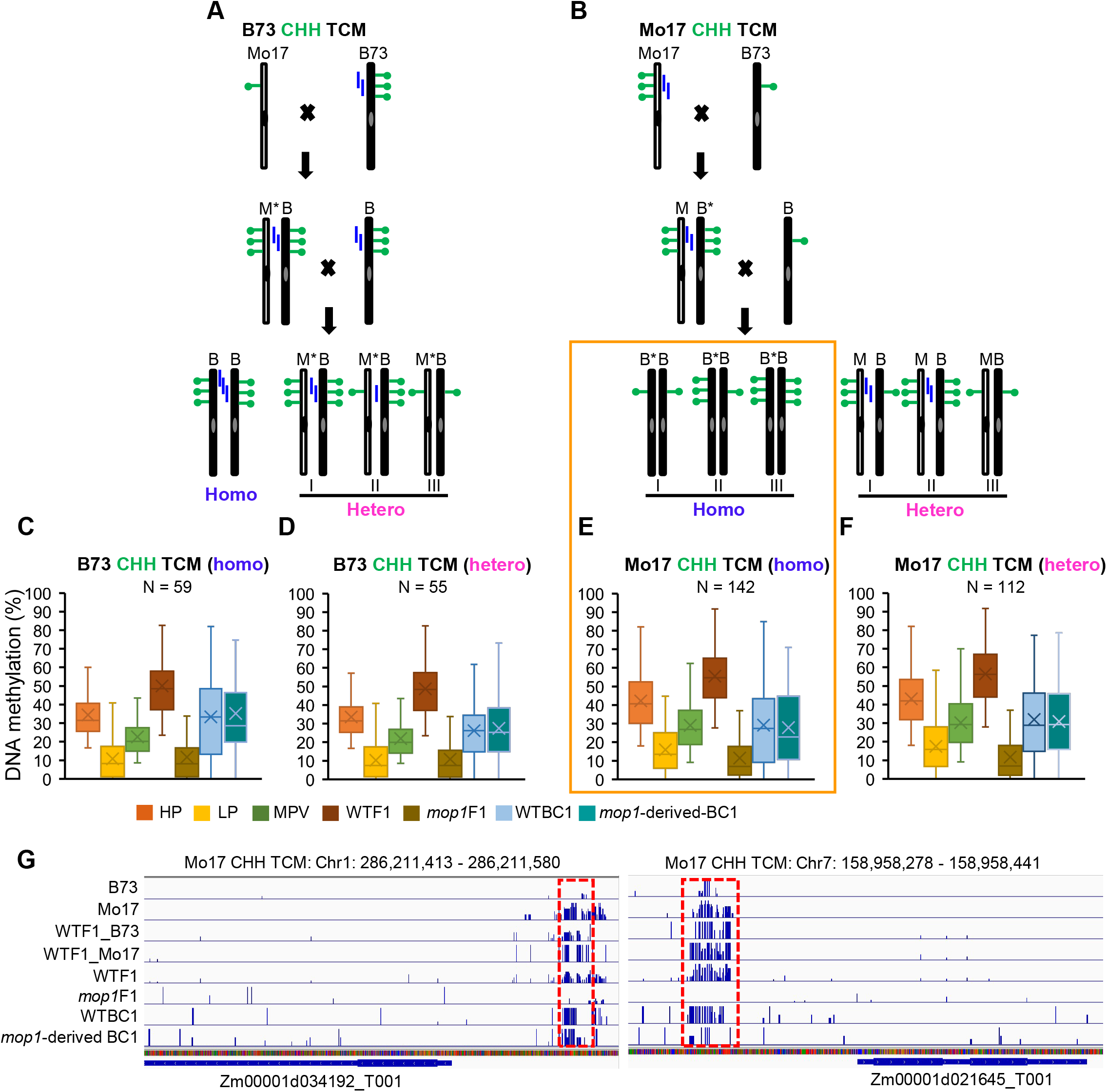
Initiation of the changes in the epigenetic state of *trans*-chromosomal CHH methylation in maize does not require Mop1. (**A**) Hypothetical model of maintenance of B73 CHH TCM DMRs. Asterisks denote the newly converted (methylated) allele. **(B)** Hypothetical model of maintenance of Mo17 CHH TCM DMRs. **(C)** Methylation changes of homozygous B73 CHH TCM DMRs. **(D)** Methylation changes of heterozygous B73 CHH TCM DMRs. **(E)** Methylation changes of homozygous Mo17 CHH TCM DMRs. **(F)** Methylation changes of heterozygous Mo17 CHH TCM DMRs. **(G)** Distribution and methylation levels of two examples of Mo17 CHH TCM DMRs. DMRs, differentially methylated regions. TCM, *trans*-chromosomal methylation. Homo, homozygous. Hetero, heterozygous. WTBC1, Mo17/B73;+/+ × B73. mop1-derived BC1, Mo17/B73;*mop1/mop1* × B73.

With respect to heritability, the Mo17 TCM DMRs (Fig 7B and Fig 8B) are more informative. In these cases, the B73 allele has become hypermethylated due to an interaction with the Mo17 allele in the F1. The BC1 plants could be either homozygous for B73, in which case one epiallele would remain more methylated if the change at B73 in the F1 were heritable, or heterozygous for Mo17 and B73, in which the B73 allele from the backcrossed parent would be expected to be newly converted to a more methylated state due to interaction with the Mo17 allele in the BC1 generation. Thus, if the change in the B73 in the F1 is heritable, we would expect to see methylation similar to the MPV of the newly converted B73 epiallele and the native B73 epiallele in the homozygotes. In the heterozygotes, we would expect a similar average level of methylation as was observed in the F1 (Fig 7B).

For CG and CHG methylation at the Mo17 TCM DMRs, the homozygotes were similar to the MPV of B73 and Mo17, rather than B73 (Fig 7E, S15E, and S15F Fig), suggesting that changes caused in the F1 plants at B73 were heritably transmitted to the next generation. In contrast, the heterozygotes, which should resemble the F1s because they carried both B73 and Mo17, were also at the MPV (Fig 7F, S15E, and S15F Fig). This suggests that the increase in methylation in B73 due to the presence of Mo17 that we observed in the F1 did not occur in BC1, again suggesting the effects we observed in F1 are not simply due to allelic interactions. For CHH methylation at the Mo17 TCM DMRs, both the homozygous and heterozygous BC1 plants had methylation at the MPV, rather than the elevated methylation observed in the F1 (Fig 8E, Fig 8F and S16D Fig). This indicated that CHH methylation that was added to the B73 allele was lost in BC1, and the presence of the Mo17 allele did not trigger methylation in B73 in the BC1 generation.

To determine whether the initiation of TCM requires the presence of MOP1, we also looked at BC1 derived from *mop1*F1 mutants (Mo17/B73;*mop1/mop1* × B73). Our expectation was that if the transfer of heritable methylation requires MOP1, backcrossed progeny of *mop1*F1 would not carry that methylation if they only carried the modified allele. The most informative class was the B73 homozygous progeny of Mo17 TCM DMRs that had shown evidence of heritable changes in methylation of the B73 allele (Fig 7E and Fig 8E). For CG and CHG TCM methylation, we find that although the *mop1* mutant had a minor effect on methylation in the F1, it had no effect on the heritability of CG and CHG methylation that had been added to the B73 allele in F1 Mo17 TCM DMRs (Fig 7E). In contrast, the substantial additional CHH methylation that was added in the F1 generation was not transmitted to the next generation of either wild type or *mop1* mutant hybrids. However, wild type progeny of *mop1*F1 mutant plants that had been nearly devoid of CHH methylation were competent to reestablish methylation (Fig 8E, orange box), suggesting that the epigenetic states at those alleles retained enough information for methylation to be targeted back to them in the wild type BC1 progeny.

Together these data suggest that methylation or demethylation triggered by hybridization can be maintained in the next generation, but that heritability varies depending on the sequence context of the methylated cytosines, and whether new methylation or demethylation is being transmitted. They also show that the elevated methylation we observed in F1 heterozygotes is likely a result of hybridization, rather than simple interaction between alleles.

## Discussion

In this study, we used hybrids as a model system to understand the initiation and maintenance of DNA methylation in maize, with a special focus on CHH methylation, which is abundant in plants, but whose functions are still poorly understood. Our analyses revealed that CHH methylation had some unique features compared to CG and CHG methylation in maize. First, only the level of CHH methylation was increased globally upon hybridization (Fig 1B). This methylation is largely enriched near genes as mCHH islands, which means that most parental CHH DMRs are located within 2 kb flanking regions of genes (Fig 2E). Second, both the high– and low-parent alleles of CHH TCM DMRs gained methylation in F1 hybrids, while only the low-parent allele gained methylation in CG and CHG TCM DMRs (Fig 3C and 3D). Furthermore, although the *mop1* mutation reduced CHH methylation globally (Fig 1B), it had its biggest effect on the CHH TCM DMRs, and the loss of CHH methylation in *mop1* resulted in additional loss of CHG methylation in these regions (Fig 4). Next, genetic variation was significantly higher in the demethylated CHH TCdM DMRs than in the methylated CHH TCM DMRs, which was not observed for CG and CHG DMRs (Fig 5E). In addition, we also provided evidence that CHH methylation in promoter regions was associated with either the suppressed or enhanced expression of flanking genes (Fig 6). Finally, we detected an overall lower level of CHH methylation in the backcross individuals relative to F1 plants (Fig 8). This suggests that the high levels of CHH methylation at individual DMRs in F1 plants is unlikely to be a consequence of *trans* interaction between these alleles alone, and is thus more likely to be a manifestation of the global effects of hybridization.

### Initiation of the changes in the epigenetic state of targets of *trans*-chromosomal CHH methylation does not depend on RdDM

Hybridization brings together two divergent genomes into one nucleus, which can induce rapid genomic and epigenomic changes, including gain or loss of DNA fragments, alteration of expression of TEs and genes, changes in splicing cites, activation of endogenous retroviruses, and epigenetic reprogramming [30, 42, 43]. It has been hypothesized that hybridization could induce a “genomic shock” that leads to the mobilization of TEs [44]. However, evidence for this is mixed and varies between species. Most reports suggest that upregulation of TEs is not a general phenomenon but that some specific TEs may change their expression level upon hybridization, such as the upregulation of *ATHILA* in the crosses of *Arabidopsis thaliana* and *Arabidopsis arenosa* [45, 46]. In our study, we observed a genome-wide increase in CHH, but not CG or CHG methylation following hybridization (Fig 1B). Based on this result, we hypothesize that CHH methylation may buffer the global effects of hybridization on transcriptional activation of TEs near genes in maize by transferring silencing information in the form of small RNAs from one genome to the other, resulting in the dramatic increases in CHH *de novo* methylation we observed in the F1 plants.

The lack of evidence for the production of new siRNAs from the target loci suggests that in many cases, this methylation is often transient, as is evidenced by the reduced heritability of CHH, CG and CHG methylation in BC1 plants (Fig 7 and Fig 8). Our analysis of gene-adjacent TEs in wild type and mutant F1 plants reveals that the cause of the increases in CHH methylation observed in F1 hybrids varies depending on location. As has been observed previously [9], the sharp increase in CHH methylation at the proximal portion of TEs near genes that are referred to as mCHH islands are dramatically reduced in the *mop1* mutant (Fig 1C and 1D). The significant increase in CHH methylation in these regions observed in F1 wild type plants are largely eliminated in the mutant as well, resulting in an overall level of CHH methylation in the F1 *mop1* mutants that is lower than both parents. That is not true in the body of gene-adjacent TEs, where CHH methylation actually increases in the F1 *mop1* mutant. At the distal edge of those TEs, although the methylation added in the F1 plants is lost, the preexisting methylation in the parents is not. In the region distal to the TEs, only a subset of the additional CHH added in the F1 plants is dependent on *Mop1* (Fig 1D). This pattern is characteristic of the vast majority of the chromosomes outside of the gene rich distal ends. Together, these data suggests that the global increase in CHH methylation observed in F1 hybrids varies with respect for a requirement for classical RdDM, with the large increases in CHH islands being the only region entirely dependent on it.

### Small RNAs are critical players in transient *trans*-chromosomal CHH methylation

Our results demonstrated that small RNAs play a critical role in triggering TCM and TCdM in hybrids and maintaining such interaction in the subsequent generation. An overall reduction of 24-nt siRNAs following hybridization has been documented in a number of plant species including maize [11, 18, 23]. In our analyses, we focused on 24-nt siRNAs specifically at TCM DMRs, and observed no significant difference in the abundance of 24-nt siRNAs between hybrids and the MPV of parents (Fig 5B), as has been seen in Arabidopsis [1]. A detailed look at 53 CHH TCM DMRs that had 24-nt siRNAs produced only in one parent showed that 34 (64%) of them had only siRNAs derived from the initially methylated parental allele, despite the fact that both alleles now had CHH methylation. Given that the precursor transcript of 24-nt siRNAs is produced by Pol IV [4, 5], this observation suggests that Pol IV in these F1 plants is only active at one of the two methylated alleles. It is unclear as to why Pol IV does not appear to recognize the newly methylated allele.

Our data also showed that CHH TCdM DMRs had significantly higher genetic variation than TCM DMRs, as has been notedly previously [1]. Given that RdDM relies on similarity between small RNAs and their targets, this may explain the reduction of methylation at TCdM DMRs. Small RNAs from the high-parent allele may be too divergent to target the low-parent allele to trigger methylation. However, it is unclear why the methylation of the high-parent allele is also reduced in the TCdM DMRs. One hypothesis that has been suggested is that small RNAs from the high-parent allele can interact with the low-parent allele unproductively, which dilutes siRNA concentration at the donor allele, which in turn weakens the methylation of the donor allele [1]. However, we do not believe this is the general mechanism for TCdM, as in our study 41.3% of these TCdM DMRs do not have any polymorphisms (Fig 3). It has been proposed that TCdM may be regulated by distal factors [47]. These distal factors have also been used to explain newly induced DMRs. In this so called ‘TCM proximity model’, a gain of methylation at TCM DMRs during hybridization spreads into flanking regions, resulting in the increased methylation in F1 at those regions, in which parental alleles have the similar methylation state [47]. However, we tested this model in our data set and did not find evidence supporting this hypothesis.

### *de novo* CHH methylation is associated with both increased and decreased expression of flanking genes

It has been proposed that mCHH islands in maize are the boundaries between highly deep heterochromatin and more active euchromatin to reinforce silencing of TEs located near genes rather than to protect the euchromatic state of the genes [9, 32, 48]. Our study is an ideal model to test this hypothesis because we can examine the effects of presence and absence of mCHH islands on the expression of the same gene in *cis* and in *trans*. For example, as shown in Fig 6A, a gene in the parent B73 does not have CHH methylation in the promoter region, but obtains methylation after hybridization, which we hypothesize is triggered in *trans* by small RNAs generated from the Mo17 allele. We demonstrated that out of the 47 CHH TCM DMRs in Mo17 (Mo17 mCHH islands), 16 (34%) were associated with suppressed gene expression, and 31 (66%) were associated with enhanced gene expression (Fig 6B), indicating that a gain of CHH methylation in their promoter regions may actually enhance their expression. Alternatively, it may be that gene properties and chromatin states may dictate the relationship between CHH islands and their flanking genes. The 31 genes whose expression appeared to be promoted by CHH methylation were generally longer, expressed higher, and with more gene body methylation than the 16 genes that seemed to be suppressed by CHH methylation (Fig 6D). This data suggests that more active genes tend to harbor “positive mCHH islands”, and lowly expressed genes more likely have “negative mCHH islands”, which were significantly enriched for the repressive histone mark H3K27me3 and were depleted with the active mark H3K4me3 (Fig 6C). We hypothesize that it is the repressive histone mark in the promoter regions suppresses the expression of flanking genes rather than the mCHH islands. Given this assumption, removal of these islands would not have significant effects on flanking gene expression. However, removal of DNA methylation may result in increase of H3K27me3 given that the activity of Polycomb-repressive complex 2, which is involved in catalyzing H3K27me3, is generally anti-correlated with DNA methylation, and likely functions after DNA demethylation [49, 50]. This probably explains why we observed these 16 genes with “negative mCHH islands” significantly reduced their expression in *mop1* mutants (Fig 6F). In contrast, the expression of the 31 genes with “positive mCHH islands” were upregulated in the *mop1* mutant although not significantly (Fig 6F), which supports the hypothesis that mCHH islands do not prevent the spread of heterochromatin silencing of genes [9]. Rather, these “positive mCHH islands” act as a border to prevent the spread of euchromatin into flanking sequences because loss of the mCHH islands in the *mop1* mutant is accompanied by additional loss of CG and CHG methylation (Fig 6E) [9].

## Materials and methods

### Genetic material construction and tissue collection

The *mop1* heterozygous plants in the Mo17 background were crossed with the *mop1* heterozygous plants in the B73 background (Mo17;*mop1-1/+* × B73;*mop1-1/+*) to generate F1 hybrid *mop1* mutants (Mo17/B73;*mop1/mop1*) and their hybrid wildtype siblings (Mo17/B73;*+/+*) (Fig 1A). The *mop1* mutation was introgressed into the B73 and Mo17 backgrounds for at least seven generations. The F1 plants and the two parental lines (B73 and Mo17) were grown in the Ecology Research Center at Miami University (Oxford, Ohio), and 5-7 cm immature ears were collected for the subsequent whole genome bisulfite sequencing (WGBS), RNA sequencing, and small RNA sequencing with two biological replicates.

### Analysis of WGBS data

DNA was isolated from the 5-7 cm immature ears of the two parents (B73 and Mo17), WTF1 (Mo17/B73;*+/+*), and *mop1*F1 (Mo17/B73;*mop1/mop1*) using the modified CTAB method. The quality of DNA was examined by Nanodrop. Library construction and subsequent WGBS were performed by Novogene. The raw reads were quality controlled by FastQC. The remaining clean reads from B73, WTF1, and *mop1*F1 were mapped to the B73 reference genome (v4) using Bismark under following parameters (-n 2, –I 50, –N 1) [51, 52]. The clean reads from the Mo17 plants were aligned against the SNP replaced Mo17 genome sequences, which were generated by taking the B73 reference sequences and replacing the nucleotides where a SNP identified by the maize Hapmap3 project was present between the two inbreds [53]. Given that treatment of DNA with bisulfite converts cytosine residues to uracil, but leaves 5-methylcytosine residues unaffected, SNPs of C to T and G to A (B73 to Mo17) were excluded from the analysis when considering the B73 allele, and SNPs of T to C and A to G (B73 to Mo17) were excluded from the analysis when considering the Mo17 allele [54]. We kept reads with perfect and unique matches for the two parents, and allowed one mismatch for the hybrids. PCR duplicates were removed using Picardtools. Additional packages including Bismark methylation extractor, bismark2bedGraph and coverage2cytosine under Bismark were used to extract the methylated cytosines, and to count methylated and unmethylated reads [55, 56].

### Identification of DMRs between parents

To identify DMRs between parents, we first filtered out the cytosines with less than three mapped reads [57]. Next, the methylation level of each cytosine was determined by the number of methylated reads out of the total number of reads covering the cytosine [58, 59]. The software ‘metilene’ was used for DMR calling between the two parents B73 and Mo17 [60]. Specially, a DMR was determined as containing at least eight cytosine sites with the distance of two adjacent cytosine sites <300 bp, and with the average methylation differences in CG and CHG >0.4 and in CHH >0.2 between the two parents [57]. These DMRs were furthered filtered by the estimated false discovery rates (FDRs) using the Benjamini-Hochberg method [1]. We only kept FDRs <0.01 for CG and CHG DMRs, and <0.05 for CHH DMRs [57].

### Determination of TCM and TCdM in WTF1

To determine the methylation patterns of the parental DMRs in WTF1, we first calculated the methylation levels at the parental DMRs in WTF1. Only DMRs with available data in all the samples were included in the analysis. Fisher’s exact test was used to compare the methylation levels of WTF1 to the MPV (middle parent value, the average methylation of the two parents), and the estimated FDRs were generated to adjust *P* values using the Benjamini-Hochberg method [1]. DMRs with an FDR <0.05 between WTF1 and the MPV were retained as significantly changed DMRs during hybridization. These DMRs were classified into TCM, which has a significantly higher level of methylation in WTF1 than the MPV, and TCdM, which has a significantly lower level of methylation in WTF1.

To further determine whether the TCM and TCdM were affected by *mop1* mutation, we first calculated the methylation levels of *mop1*F1 at TCM and TCdM DMRs. For CG and CHG TCM and TCdM, DMRs with the changes in methylation levels between *mop1*F1 and WTF1 <-0.4 or >0.4 were considered as significantly affected by the mutation. For CHH TCM and TCdM, DMRs with the changes in methylation levels <-0.2 or >0.2 were considered as significantly changed in the mutants.

### Identification of the newly induced DMRs in WTF1

To identify the newly induced DMRs in WTF1 that are not differentially methylated in the parents, we used mpileup in the samtools package and SNPs between B73 and Mo17 to obtain the allele-specific reads from WTF1 [61]. Next, these allele-specific reads were used to calculate methylation levels as described above, and ‘metilene’ was used for DMR detection between the two alleles in WTF1 [60]. The same cutoffs are used for defining new DMRs as for the detection of parental DMRs. The methylation levels of these newly induced DMRs were further compared with the methylation levels of the two parents using Fisher’s exact test (FDR <0.05). The DMRs that have similar methylation levels between the two parents but exhibit significantly different methylation levels between the two alleles in WTF1 were defined as new DMRs. The illustration is shown in A and S12B Fig.

### Analysis of the inheritance of TCM and TCdM in BC1

We backcrossed WTF1 (Mo17/B73;*+/+*) and *mop1*F1 (Mo17/B73;*mop1/mop1*) with B73 (Mo17/B73;*+/+* × B73 and Mo17/B73;*mop1/mop1* × B73) to generate the BC1 generation. We collected 5-7 cm immature ears from eight WTBC1 plants and eight *mop1*-derived-BC1 plants for WGBS (Fig 1A). The methylation analysis for BC1 is the same as that for parents and WTF1. Next, we compared the methylation levels at the TCM and TCdM DMRs among WTBC1, *mop1*– derived-BC1, WTF1, *mop1*F1, and parents. The “intersect” function in BEDTools was used to access all the cytosines in BC1 that are at the TCM and TCdM DMRs, and these cytosines were used to calculate the average methylation levels across all the BC1 individuals in those regions. As shown in Fig 8A,B, because we only sequenced the BC1 individuals derived from the backcrosses of F1 and B73, we separated B73/B73 homozygous and B73/Mo17 heterozygous genotypes at each TCM in BC1 using samtools mpileup and the SNPs between B73 and Mo17 [53, 61], same as what we did for the determination of allele specific reads in F1.

### Distribution analysis of DMRs in different genomic locations

To classify the DMRs in different genomic locations, we compared the locations of the DMRs with gene and transposable element annotations, which were downloaded from MaizeGDB, https://www.maizegdb.org/, using intersect function in BEDTools [62]. If one DMR is dropped to two different types of annotation, we followed the order gene bodies, 2 kb upstream of genes, 2kb downstream of genes, TEs, and unclassified regions. DMRs in the 2 kb up and downstream regions of genes were further separated into those with and without TEs depending on whether there is a TE insertion in the 2 kb flanking regions.

### Analysis of small RNA-seq data

The same genetic materials for B73 and Mo17, WTF1, and *mop1*F1 were used for small RNA sequencing with two biological replicates. The raw reads were quality controlled by FastQC. The clean reads were aligned against the Rfam database (v14.6) to remove rRNAs, tRNAs, snRNAs, and snoRNAs [56]. The remaining reads with the length of 18-26 nt were retained for further mapping to the genomes. The reads from B73, WTF1, and *mop1*F1 were mapped to the B73 reference genome (v4), and the reads from Mo17 were mapped to the SNP replaced Mo17 genome sequences using bowtie [52, 63], as was done for our methylation analysis. For the parents, only perfectly and uniquely mapped reads were kept, and one mismatch was allowed for the F1 hybrids. The small RNA values were adjusted to total abundance of all mature microRNAs following the previous research to remove the artificial increase of 22-nt siRNAs in *mop1* mutants caused by normalization [35]. The intersect module in BEDTools was used to compare the mapping results (sam files) to the positions of DMRs to obtain the 24-nt small RNA reads that are in the DMRs [62]. These 24-nt small RNAs were used to calculate the expression of small RNAs of the DMRs. To access allele specific small RNA expression, samtools mpileup and SNPs at small RNAs between B73 and Mo17 were used [61].

### Analysis of mRNA-seq data

The mRNA from the same genetic materials were sequenced with two biological replicates. The raw reads were quality controlled by FastQC, and the low-quality reads and the adapter sequences were removed by Trimmomatic [64]. We mapped the cleaned reads of B73, WTF1, and *mop1*F1 to the B73 reference genome (v4) [52], and the reads from Mo17 to the SNP replaced Mo17 genome that was generated by replacing the B73 genome with the SNPs between Mo17 and B73 using Hisat2 with one mismatch [65]. Next, HTSeq-count was used to calculate the total number of reads of each gene [66]. These values were loaded to DESeq2 to identify genes that were differentially expressed between WTF1 and parents, and between WTF1 and *mop1*F1 [67]. To determine allele specific expression of each gene in F1, the mpileup function in samtools and SNPs between B73 and Mo17 were used to access allele specific reads [61], which were further used in DESeq2 to identify differential expression of the two alleles [67].

### Accession Numbers

The raw and processed data of whole genome bisulfite, mRNA and small RNA sequencing presented in this study have been deposited in NCBI Gene Expression Omnibus under the accession number GSE222155.

### Supporting information

**S1 Fig. Whole genome levels of DNA methylation among parents, hybrids and mutants.** The average methylation of the overall cytosine (total C), CG, CHG, and CHH on the whole genome in parents, WTF1, and *mop1*F1.

**S2 Fig. CHH methylation is globally increased in hybrids.** Nine of the 10 maize chromosomes are shown here. Methylation levels were measured in 1 Mb windows with 500 kb shift. Here WTF1 indicates the sibling of *mop1*F1. The shaded boxes represent pericentromeric region of each chromosome.

**S3 Fig. The length distribution of the DMRs identified between parents**. (**A**) CG DMRs. (**B**) CHG DMRs. **(C)** CHH DMRs. DMRs, differentially methylated regions.

**S4 Fig. Genomic distribution of unchanged (NC) parental DMRs. (A)** CG DMRs. **(B)** CHG DMRs. **(C)** CHH DMRs. **(D)** The types of TEs at the categories of 2 kb upstream of genes with TEs and 2 kb downstream of genes with TEs **A-C**. 2 kb upstream of genes with TEs (transposable elements) and 2 kb downstream of genes with TEs indicate both the DMRs and TEs are located within the 2 kb of genes.

**S5 Fig. Methylation changes at the unchanged (NC) DMRs**. HP, high parent (parent with higher methylation). HA, high-parent allele in F1. LP, low parent (parent with lower methylation). LA, low-parent allele in F1. Average means the average between the two parents, or between the two alleles in WTF1 and *mop1*F1. DMRs, differentially methylated regions.

**S6 Fig. Genomic distribution of *mop1*-affected DMRs. (A)** CG DMRs. **(B)** CHG DMRs. **(C)** CHH DMRs. 2 kb upstream of genes with TEs (transposable elements) and 2kb downstream of genes with TEs indicate both the DMRs and TEs are located within the 2 kb of genes. DMRs, differentially methylated regions.

**S7 Fig. The production of 24-nt small interfering RNAs (siRNAs) from gene bodies and flanking regions. (A)** Patterns of CHH methylation in and flanking genes. **(B)** The expression of 24-nt siRNAs on gene bodies and flanking regions. TSS, transcription start site. TTS, transcription termination site. TP10M = siRNA reads/total unique mapped reads *10,000,000.

**S8 Fig. The high parent has significantly more 24-nt small interfering RNAs (siRNAs).** HP, high parent (parent with higher methylation). LP, low parent (parent with lower methylation). DMRs, differentially methylated regions. RPKM, 24-nt siRNA reads per kilobase (DMR length) per million uniquely mapped reads. **, *P* < 0.01. Student’s *t* test.

**S9 Fig. Comparisons of 24-nt small interfering RNAs (siRNAs) at unchanged (NC), TCM and TCdM DMRs. (A)** NC DMRs. **(B)** TCM DMRs. **(C)** TCdM DMRs. HP, high parent (parent with higher methylation). LP, low parent (parent with lower methylation). MPV, the middle parent value. DMRs, differentially methylated regions. TCM, *trans*-chromosomal methylation. TCdM, *trans*-chromosomal demethylation. RPKM, 24-nt siRNA reads per kilobase (DMR length) per million uniquely mapped reads. **, *P* < 0.01, *, *P* < 0.05. Student’s *t* test.

**S10 Fig. Expression of genes involved in the transcriptional gene silencing pathway**.

**S11 Fig. CHH methylation is associated with both suppressed and enhanced expression of their flanking genes. (A)** Two possible scenarios of the effects of CHH methylation on gene expression. Here only shows the examples of Mo17 CHH TCM DMRs. **(B)** Expression values of the 16 and 31 genes that are associated with suppressed and enhanced expression by flanking CHH DMRs respectively between the two parents (B73 and Mo17). *, *P* < 0.05. Student’s paired *t* test. **(C)** DNA methylation levels between the 16 and 31 CHH DMRs that are with suppressed and enhanced expression of flanking genes. *, *P* < 0.05. Student’s *t* test. **(D)** Gene length including introns between the 16 and 31 genes. *, *P* < 0.05. Student’s *t* test.

**S12 Fig. Most new CG and CHG DMRs lose methylation, and most new CHH DMRs gain methylation in WTF1. (A)** and **(B)** Two hypothetical models of new CG, CHG and CHH DMRs induced in WTF1. **(C)** Comparisons of CG, CHG and CHH methylation at DMRs following the Model A. **(D)** Comparisons of CG, CHG and CHH methylation at DMRs following the Model B. HP/HA indicates high parent or high-parent allele in F1, and LP/LA represents low parent or low-parent allele in F1. Average means the average between the two parents, or between the two alleles in WTF1 and *mop1*F1. DMRs, differentially methylated regions.

**S13 Fig. Newly induced CG and CHG DMRs are largely located in transposable elements**. **(A)** and **(B)** Two hypothetical models of new CG, CHG and CHH DMRs induced in WTF1. **(C)** Number of B73 and Mo17 hyper DMRs in Model A. **(D)** Number of B73 and Mo17 hyper DMRs in Model B. **(E)** The distribution of CG, CHG and CHH DMRs in Model A. **(F)** The distribution of CG, CHG and CHH DMRs in Model B. 2 kb upstream of genes with TEs (transposable elements) and 2kb downstream of genes with TEs indicate both the DMRs and TEs are located within the 2 kb of genes.

**S14 Fig. No significant changes in small RNAs between the two parents, and between the hybrids and parents. (A)** and **(B)** Two hypothetical models of new CHH DMRs induced in WTF1. **(C)** 24-nt small interfering RNA (siRNAs) of new CHH DMRs in Model A. **(D)** 24-nt siRNAs of new CHH DMRs in Model B. **(E)** Ratios of 24 nt siRNAs of high parent to low parent, and of high-parent allele to low-parent allele at the new CHH DMRs in Model A. **(F)** Ratios of 24-nt siRNAs of high parent to low parent, and of high-parent allele to low-parent allele at the new CHH DMRs in Model B. HP, high parent (parent with higher methylation). LP, low parent (parent with lower methylation). MPV, the middle parent value. **, *P* < 0.01, Student’s *t* test. DMRs, differentially methylated regions.

**S15 Fig. Inheritance of newly triggered methylation at CG and CHG TCM DMRs in the backcrossed generation**. **(A)** Hypothetical model of maintenance of B73 CG and CHG TCM DMRs. Asterisk denotes the newly converted (methylated) allele. **(B)** Hypothetical model of maintenance of Mo17 CG and CHG TCM DMRs. **(C)** Methylation changes of B73 CG TCM DMRs. **(D)** Methylation changes of B73 CHG TCM DMRs. **(E)** Methylation changes of Mo17 CG TCM DMRs. **(F)** Methylation changes of Mo17 CHG TCM DMRs. DMRs, differentially methylated regions. TCM, *trans*-chromosomal methylation. Homo, homozygous. Hetero, heterozygous. WTBC1, Mo17/B73;+/+ × B73. *mop1*-derived BC1, Mo17/B73;*mop1/mop1* × B73.

**S16 Fig. Inheritance of newly triggered methylation at CHH TCM DMRs in the backcrossed generation**. **(A)** Hypothetical model of maintenance of B73 CHH TCM DMRs. Asterisk denotes the newly converted (methylated) allele. **(B)** Hypothetical model of maintenance of Mo17 CHH TCM DMRs. **(C)** Methylation changes of B73 CHH TCM DMRs. **(D)** Methylation changes of Mo17 CHH TCM DMRs. DMRs, differentially methylated regions. TCM, *trans*-chromosomal methylation. Homo, homozygous. Hetero, heterozygous. WTBC1, Mo17/B73;+/+ × B73. *mop1*-derived BC1, Mo17/B73;*mop1/mop1* × B73.

**S1 Table. The summary of raw reads of different samples.**

**S2 Table. The overall patterns of cytosine methylation in parents, WTF1, and mutant F1.**

**S3 Table. DMRs identified between parents (B73 and Mo17).**

**S4 Table. Number of changed DMRs in the other two cytosine contexts at the mop1– affected CG, CHG, and CHH DMRs.**

**S5 Table. Differentially expressed genes involved in the transcriptional gene silencing pathway between parents.**

**S6 Table. Differentially expressed genes involved in the transcriptional gene silencing pathway between MPV and F1.**

**S7 Table. Inheritance of CG and CHG TCM and TCdM in the backcrossed generation (BC1).**

**S8 Table. Inheritance of CHH TCM and TCdM in the backcrossed generation (BC1).**

## Acknowledgements

We are grateful to Nathan M Springer and Karen M. McGinnis for critical reading of the manuscript. We thank Ohio Supercomputer Center for providing us with the computational resources to perform the analysis.

## Author contributions

**Conceptualization:** Beibei Liu, Damon Lisch, Meixia Zhao.

**Data curation:** Beibei Liu, Diya Yang, Dafang Wang.

**Funding acquisition:** Dafang Wang, Meixia Zhao.

**Investigation:** Beibei Liu, Diya Yang, Meixia Zhao.

**Methodology:** Beibei Liu, Diya Yang, Chun Liang, Meixia Zhao.

**Project administration:** Meixia Zhao.

**Resources:** Damon Lisch, Meixia Zhao.

**Software:** Beibei Liu, Meixia Zhao.

**Supervision:** Beibei Liu, Meixia Zhao.

**Validation:** Beibei Liu, Meixia Zhao.

**Visualization:** Beibei Liu, Meixia Zhao.

**Writing – original draft:** Beibei Liu, Meixia Zhao.

**Writing – review & editing:** Beibei Liu, Dafang Wang, Chun Liang, Jianping Wang, Damon Lisch, Meixia Zhao.

