## Supplemental Figures and Tables for "Heritable changes of epialleles in maize can be triggered in the absence of DNA methylation"

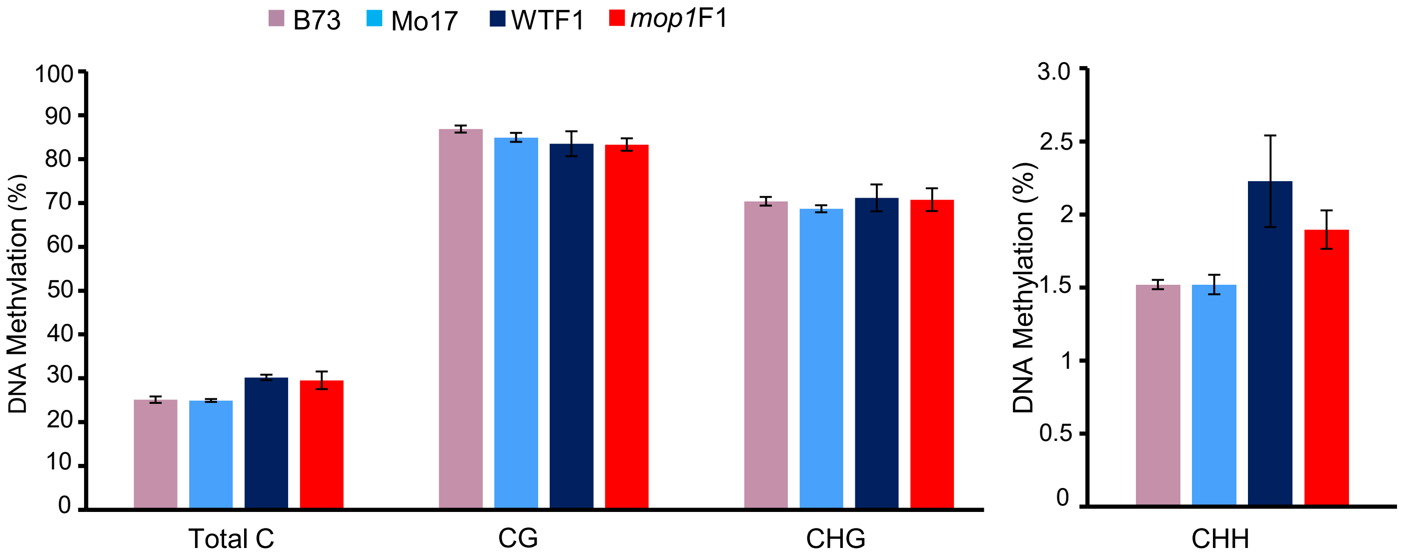


**S1 Fig. Whole genome levels of DNA methylation among parents, hybrids and mutants**.

The average methylation of the overall cytosine (total C), CG, CHG, and CHH on the whole genome in parents, WTF1, and *mop1*F1.

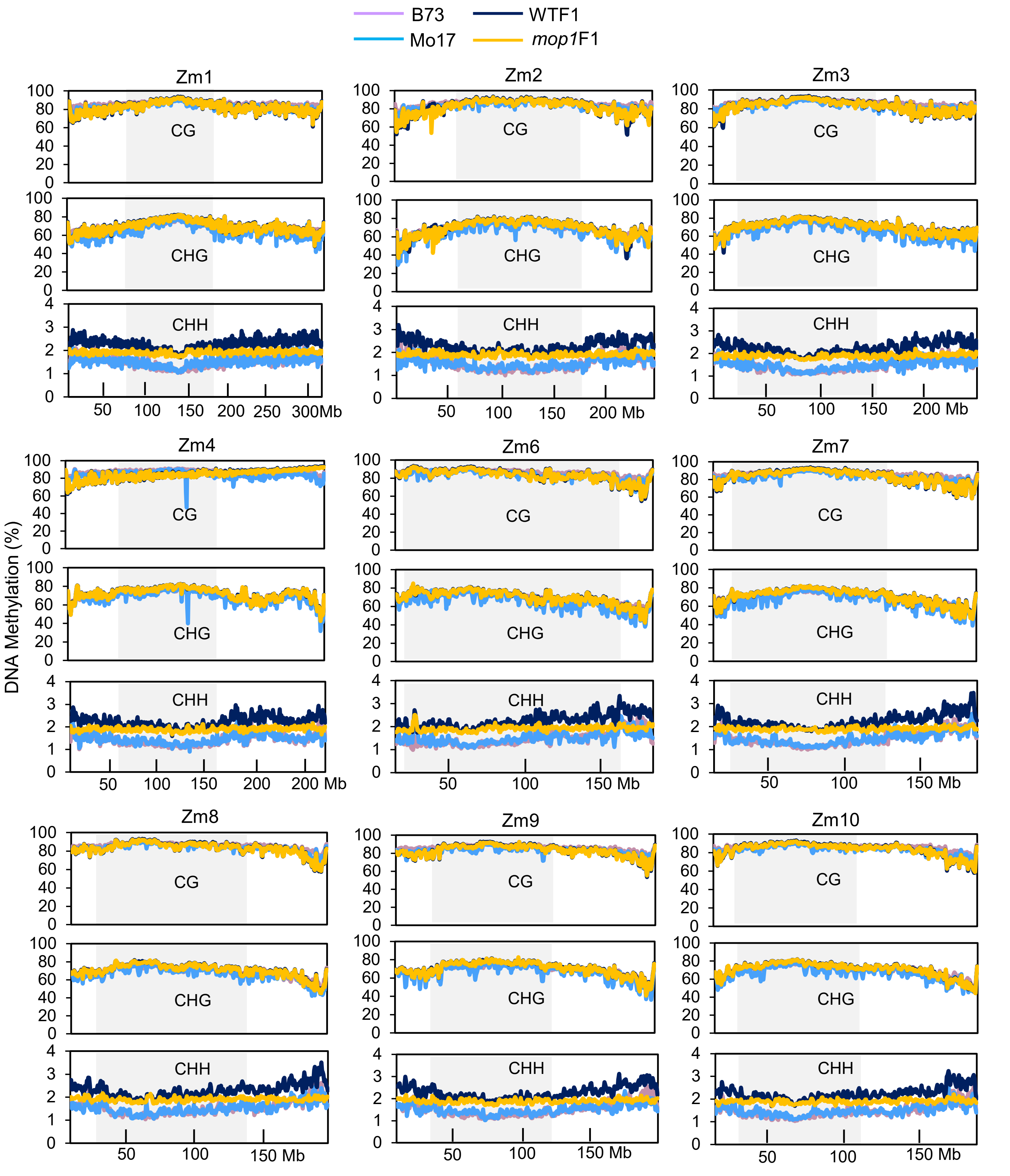


**S2 Fig. CHH methylation is globally increased in hybrids.**
Nine of the 10 maize chromosomes are shown here. Methylation levels were measured in 1 Mb windows with 500 kb shift. Here WTF1 indicates the sibling of *mop1*F1. The shaded boxes represent pericentromeric region of each chromosome.

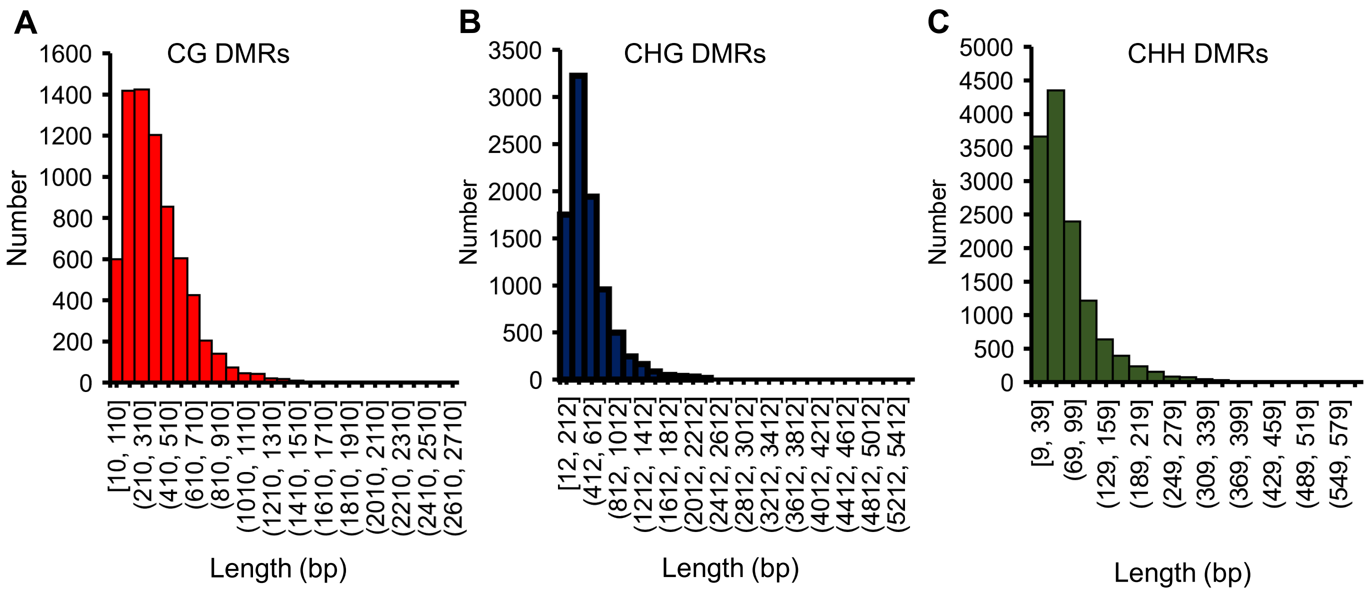


**S3 Fig. The length distribution of the DMRs identified between parents.**
**(A)** CG DMRs. **(B)** CHG DMRs. **(C)** CHH DMRs. DMRs, differentially methylated regions.

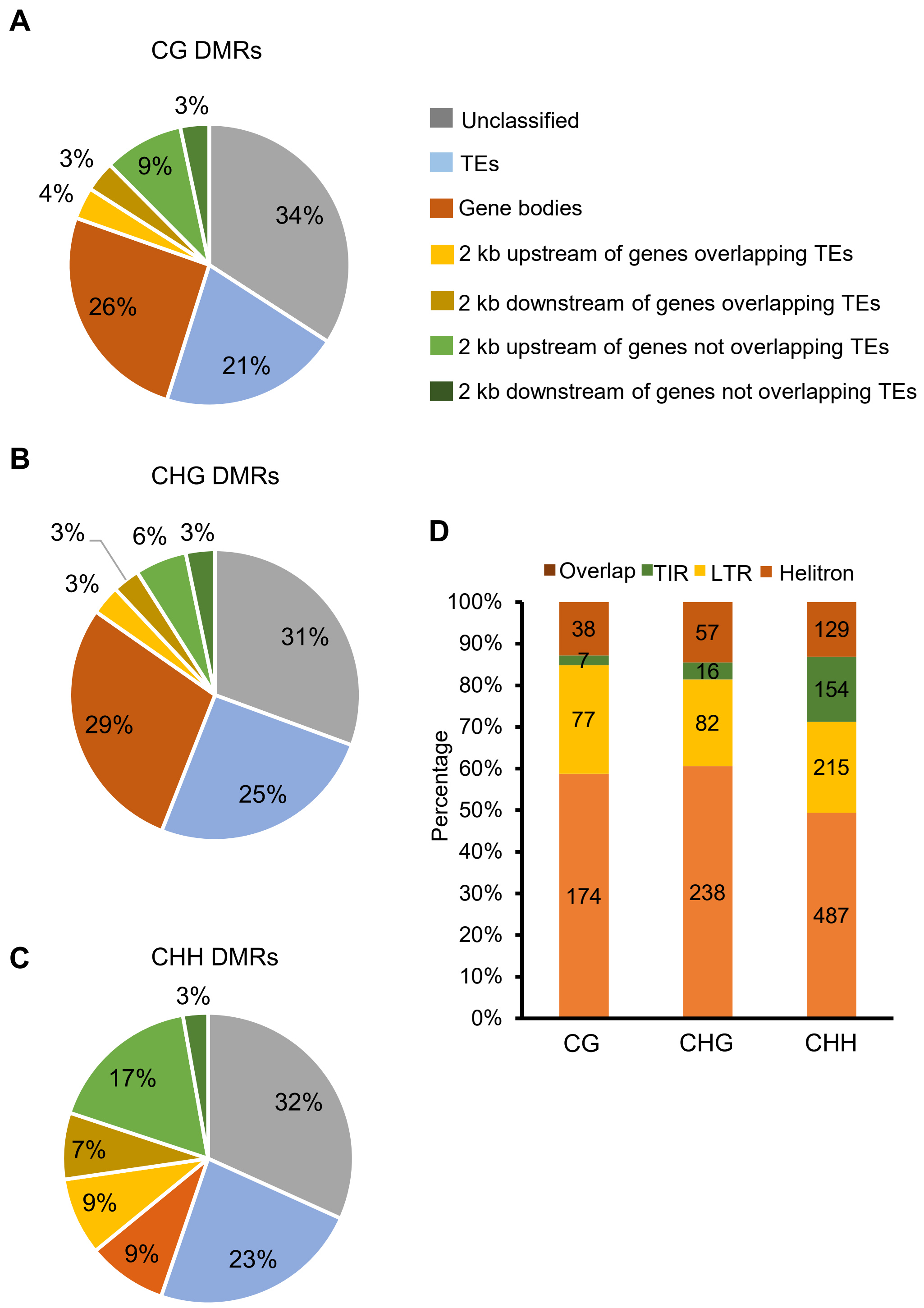


**S4 Fig. Genomic distribution of unchanged (NC) parental DMRs.**
**(A)** CG DMRs. **(B)** CHG DMRs. **(C)** CHH DMRs. **(D)** The types of TEs at the categories of 2 kb upstream of genes with TEs and 2 kb downstream of genes with TEs **A-C**. 2 kb upstream of genes with TEs (transposable elements) and 2 kb downstream of genes with TEs indicate both the DMRs and TEs are located within the 2 kb of genes.

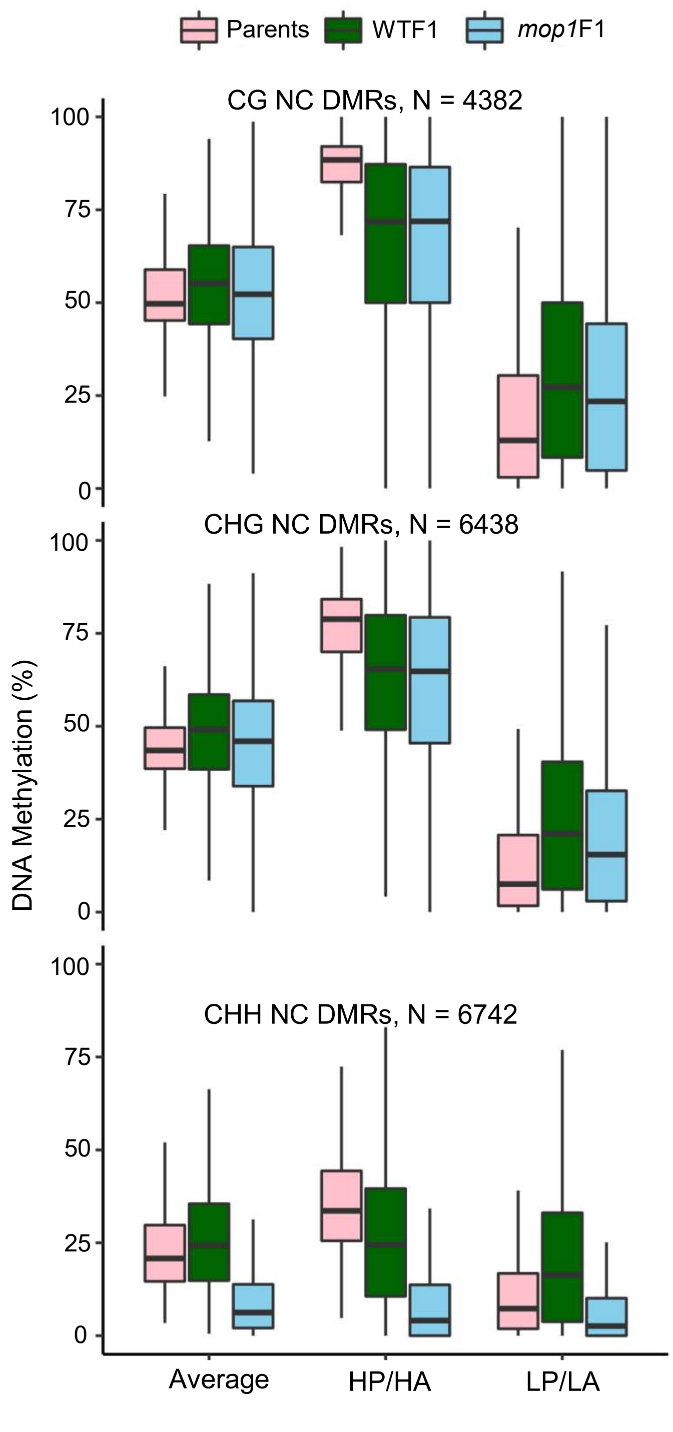


**S5 Fig. Methylation changes at the unchanged (NC) DMRs**.
HP, high parent (parent with higher methylation). HA, high-parent allele in F1. LP, low parent (parent with lower methylation). LA, low-parent allele in F1. Average means the average between the two parents, or between the two alleles in WTF1 and *mop1*F1. DMRs, differentially methylated regions.

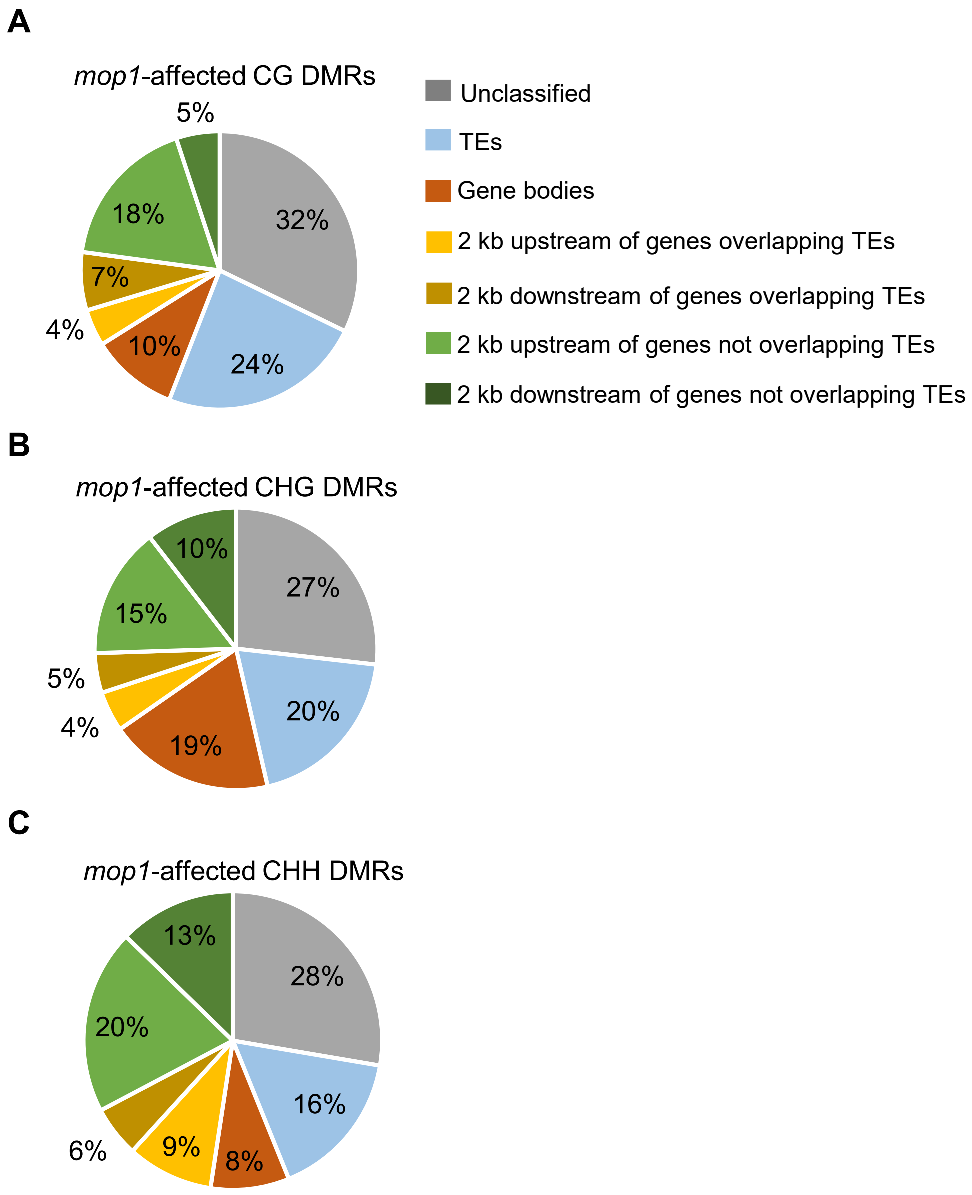


**S6 Fig. Genomic distribution of *mop1*-affected DMRs.**
**(A)** CG DMRs. **(B)** CHG DMRs. **(C)** CHH DMRs. 2 kb upstream of genes with TEs (transposable elements) and 2kb downstream of genes with TEs indicate both the DMRs and TEs are located within the 2 kb of genes. DMRs, differentially methylated regions.

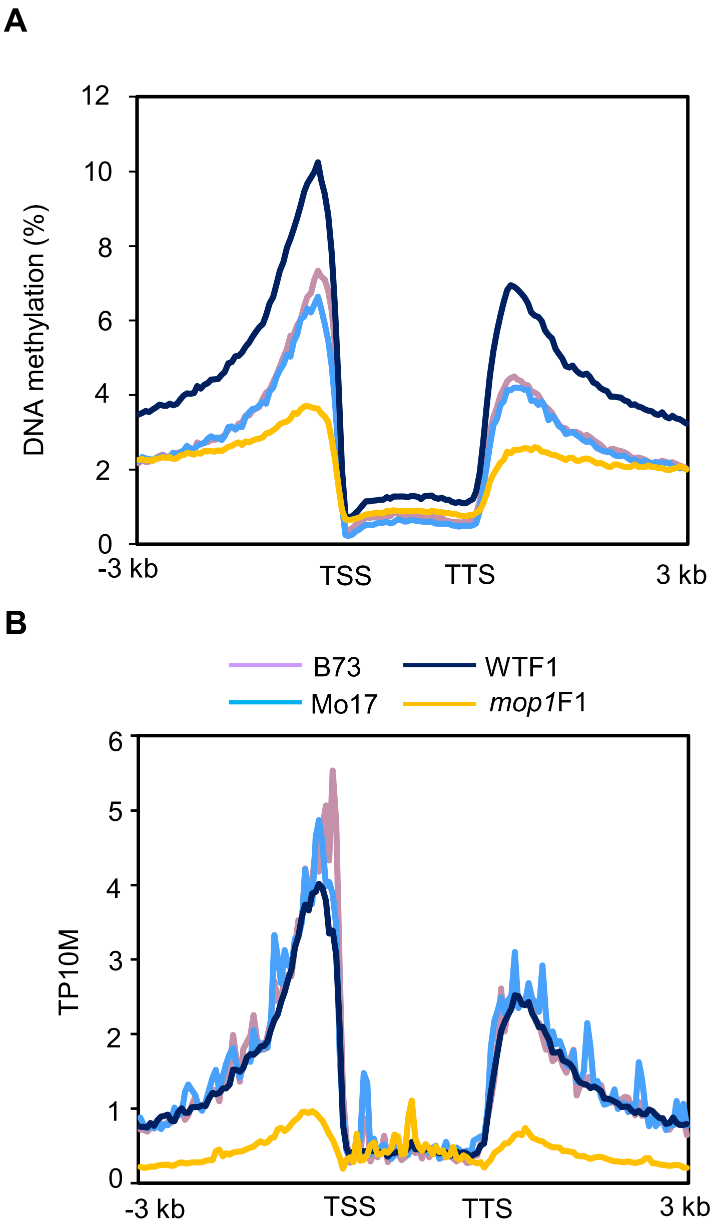


**S7 Fig. The production of 24-nt small interfering RNAs (siRNAs) from gene bodies and flanking regions.**
**(A)** Patterns of CHH methylation in and flanking genes.
**(B)** The expression of 24-nt siRNAs on gene bodies and flanking regions. TSS, transcription start site. TTS, transcription termination site. TP10M = siRNA reads/total unique mapped reads *10,000,000.

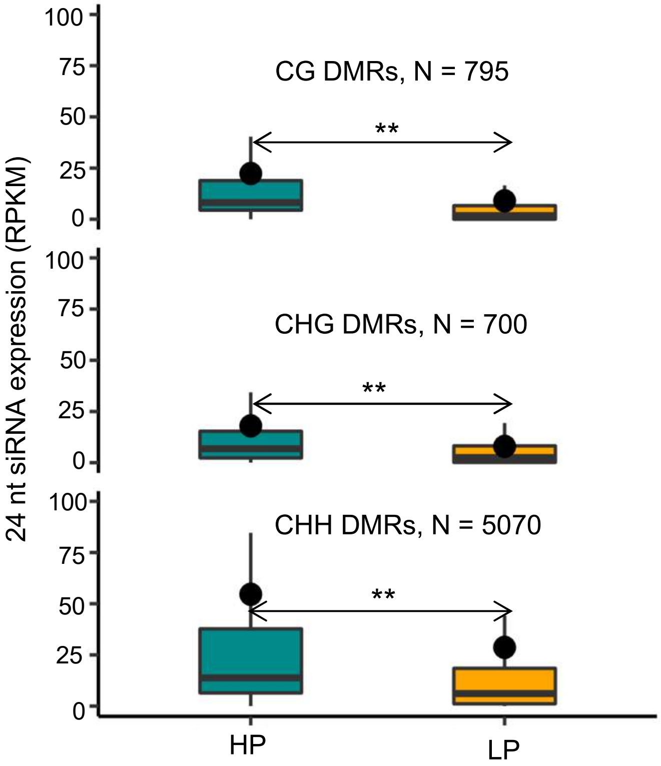


**S8 Fig. The high parent has significantly more 24-nt small interfering RNAs (siRNAs).**

HP, high parent (parent with higher methylation). LP, low parent (parent with lower methylation). DMRs, differentially methylated regions. RPKM, 24-nt siRNA reads per kilobase (DMR length) per million uniquely mapped reads. **, $P < 0.01$. Student's *t* test.

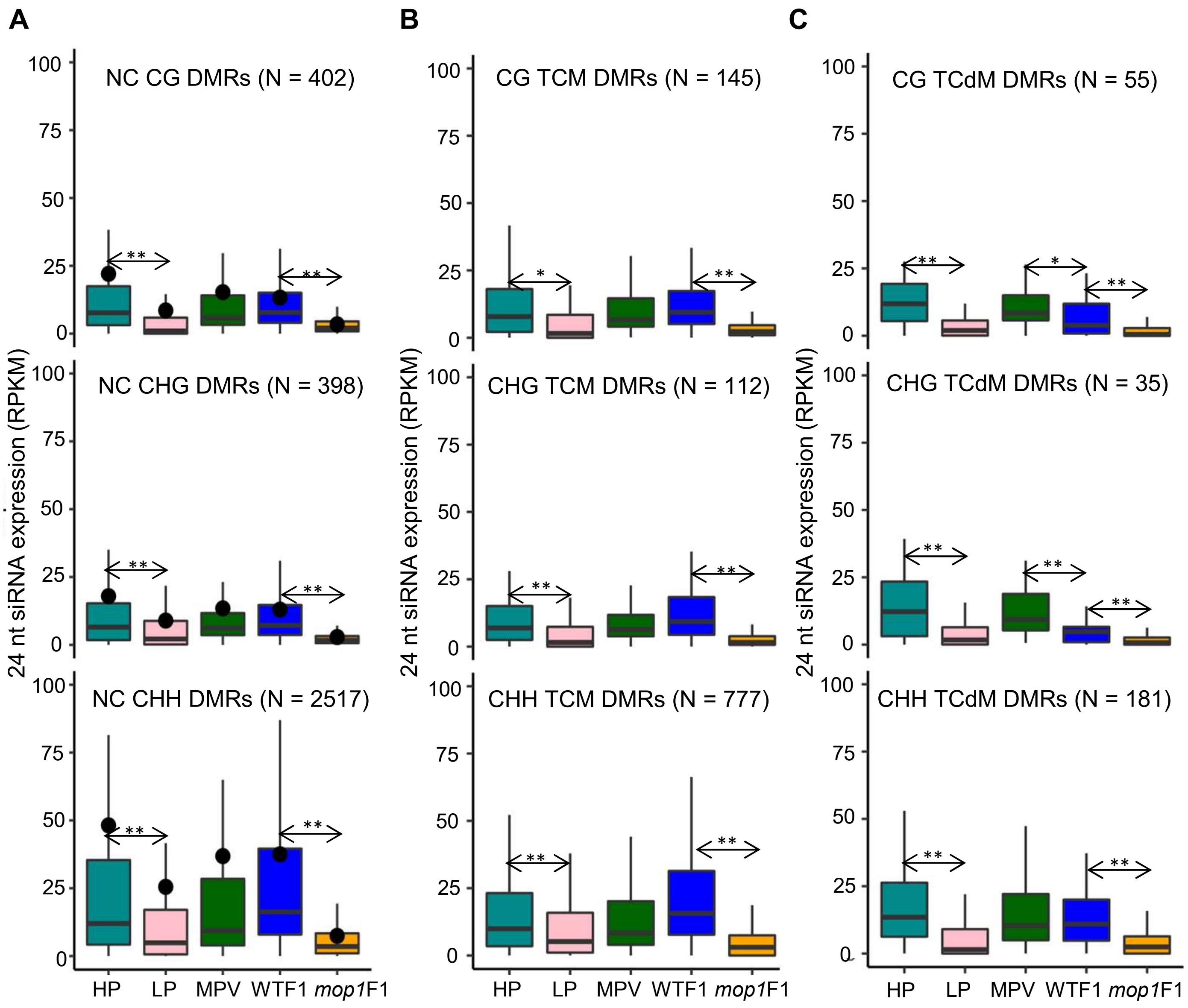


**S9 Fig. Comparisons of 24-nt small interfering RNAs (siRNAs) at unchanged (NC), TCM and TCdM DMRs.**

**(A)** NC DMRs. **(B)** TCM DMRs. **(C)** TCdM DMRs. HP, high parent (parent with higher methylation). LP, low parent (parent with lower methylation). MPV, the middle parent value. DMRs, differentially methylated regions. TCM, *trans*-chromosomal methylation. TCdM, *trans*-chromosomal demethylation. RPKM, 24-nt siRNA reads per kilobase (DMR length) per million uniquely mapped reads. **, $P < 0.01$, *, $P < 0.05$. Student's *t* test.

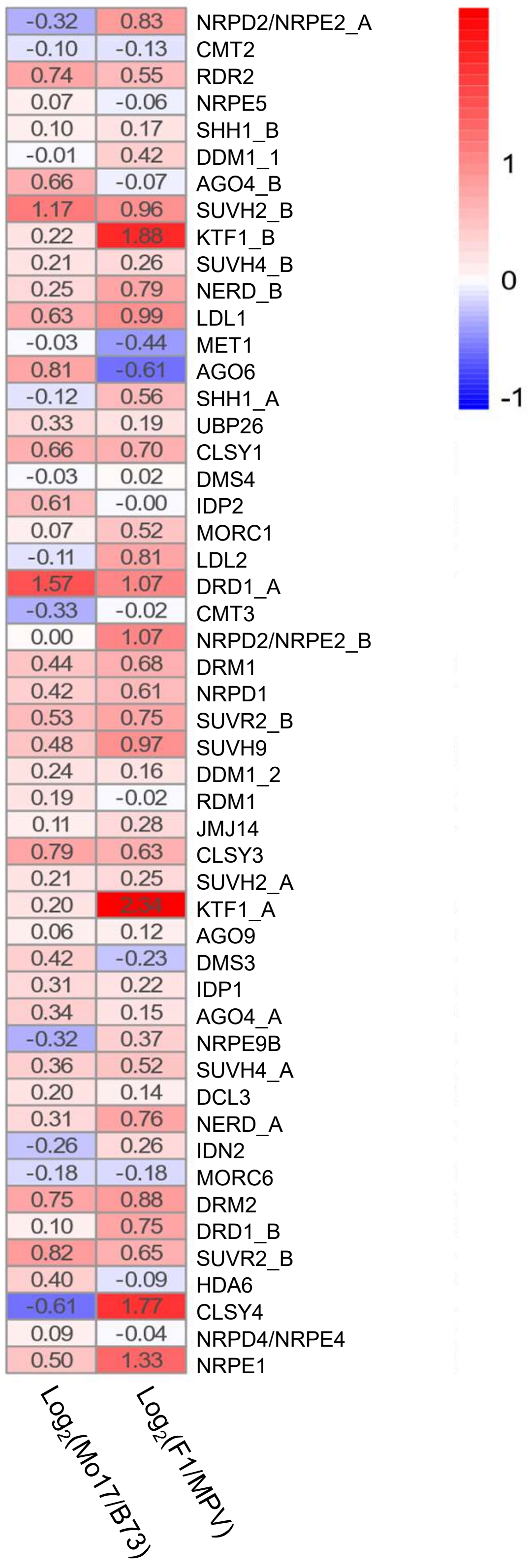


**S10 Fig. Expression of genes involved in the transcriptional gene silencing pathway.**

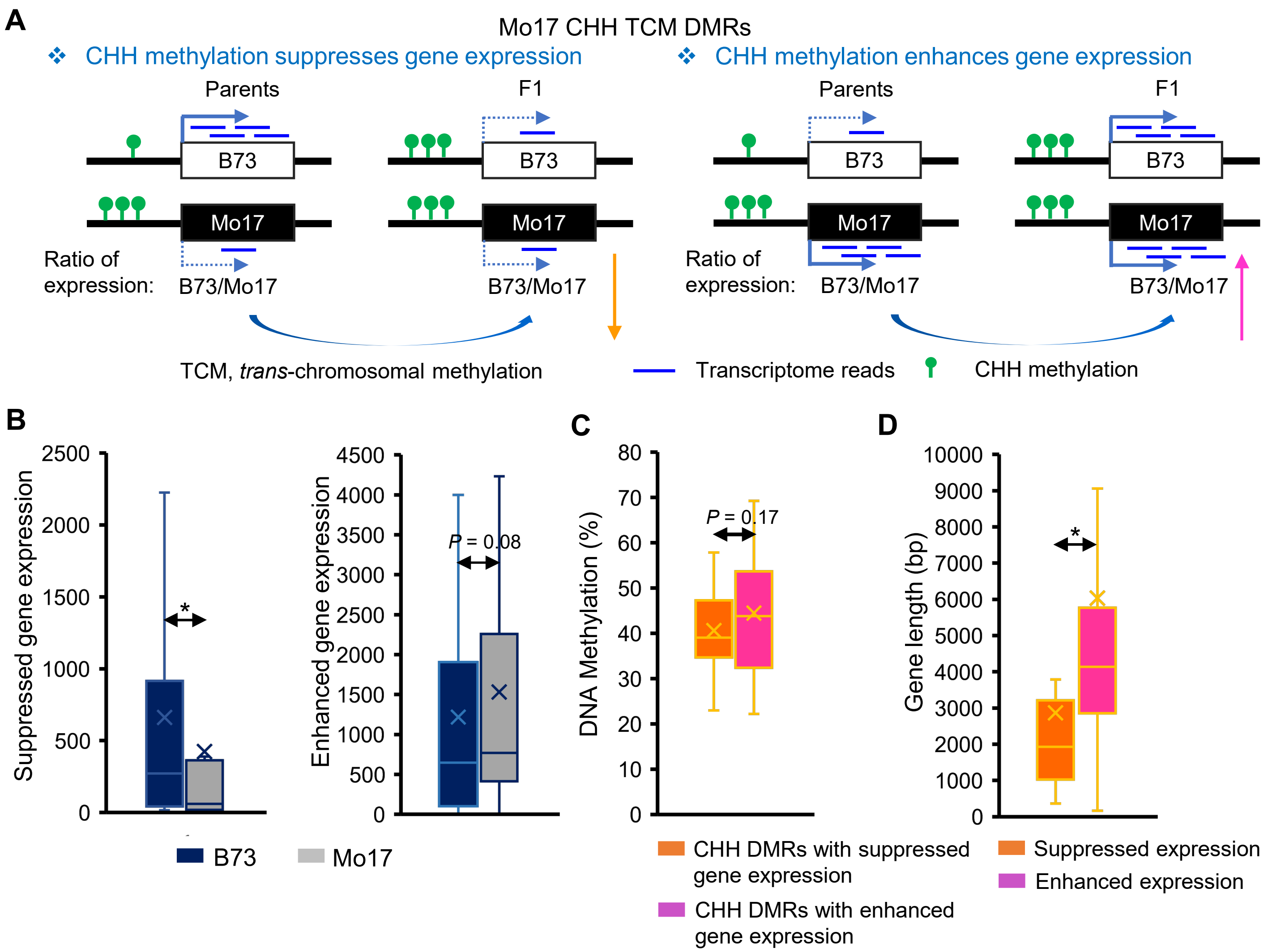


**S11 Fig. CHH methylation is associated with both suppressed and enhanced expression of their flanking genes.**

**(A)** Two possible scenarios of the effects of CHH methylation on gene expression. Here only shows the examples of Mo17 CHH TCM DMRs. **(B)** Expression values of the 16 and 31 genes that are associated with suppressed and enhanced expression by flanking CHH DMRs respectively between the two parents (B73 and Mo17). *, $P < 0.05$. Student's paired *t* test. **(C)** DNA methylation levels between the 16 and 31 CHH DMRs that are with suppressed and enhanced expression of flanking genes. *, $P < 0.05$. Student's *t* test. **(D)** Gene length including introns between the 16 and 31 genes. *, $P < 0.05$. Student's *t* test.

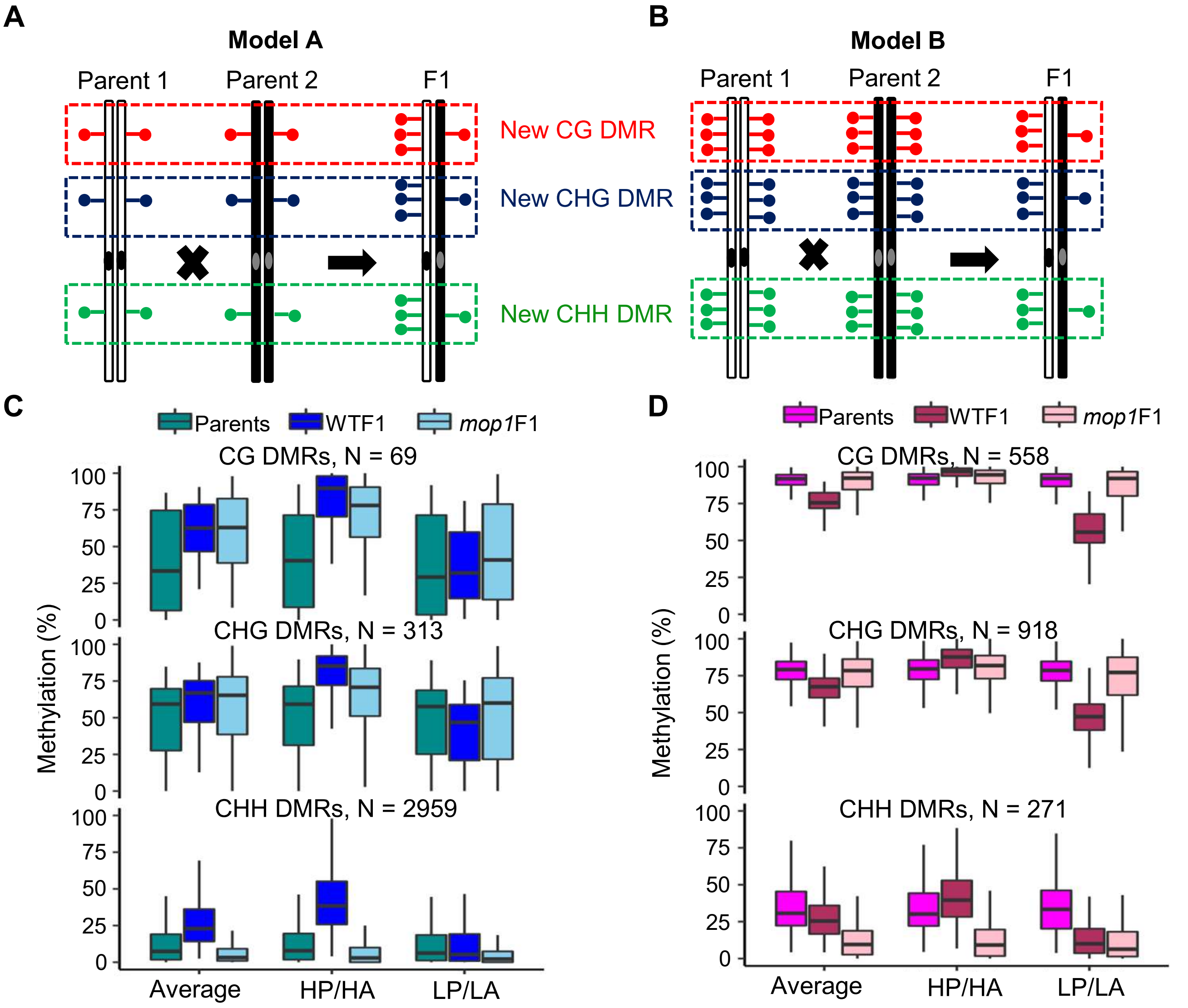


**S12 Fig. Most new CG and CHG DMRs lose methylation, and most new CHH DMRs gain methylation in WTF1.**

**(A)** and **(B)** Two hypothetical models of new CG, CHG and CHH DMRs induced in WTF1. **(C)** Comparisons of CG, CHG and CHH methylation at DMRs following the Model A. **(D)** Comparisons of CG, CHG and CHH methylation at DMRs following the Model B. HP/HA indicates high parent or high-parent allele in F1, and LP/LA represents low parent or low-parent allele in F1. Average means the average between the two parents, or between the two alleles in WTF1 and *mop1*F1. DMRs, differentially methylated regions.

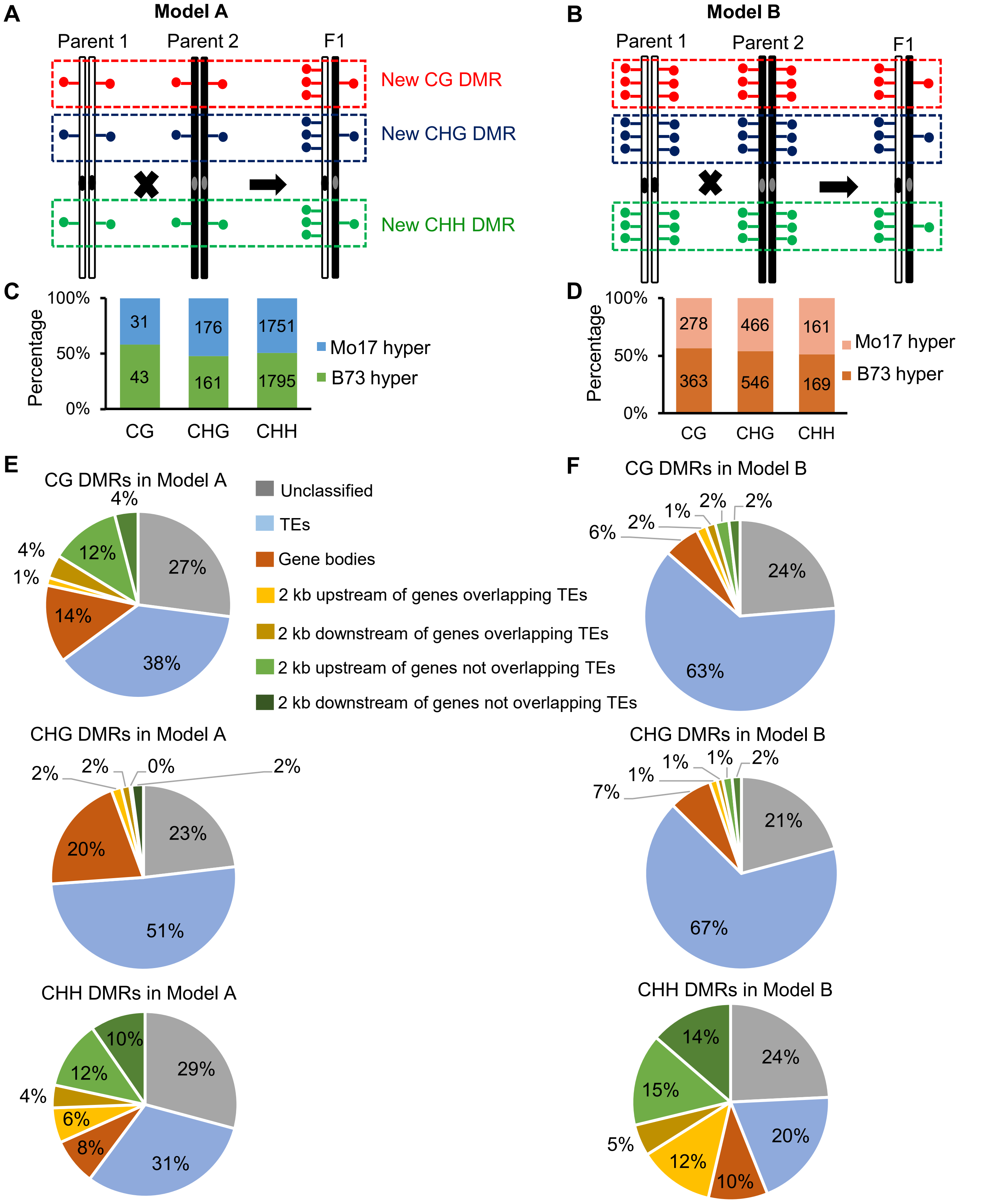


**S13 Fig. Newly induced CG and CHG DMRs are largely located in transposable elements.**

**(A)** and **(B)** Two hypothetical models of new CG, CHG and CHH DMRs induced in WTF1. **(C)** Number of B73 and Mo17 hyper DMRs in Model A. **(D)** Number of B73 and Mo17 hyper DMRs in Model B. **(E)** The distribution of CG, CHG and CHH DMRs in Model A. **(F)** The distribution of CG, CHG and CHH DMRs in Model B. 2 kb upstream of genes with TEs (transposable elements) and 2kb downstream of genes with TEs indicate both the DMRs and TEs are located within the 2 kb of genes.

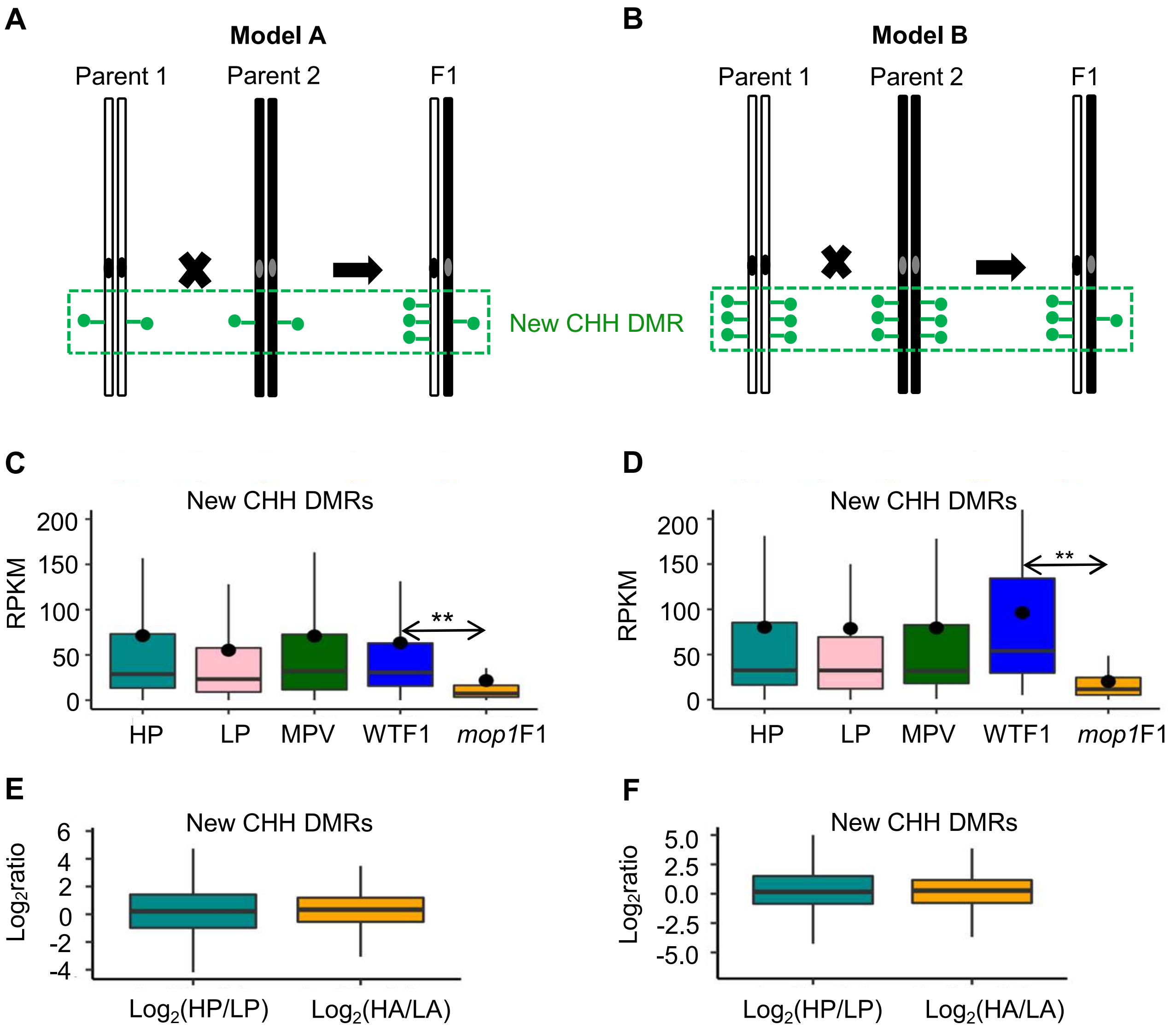


**S14 Fig. No significant changes in small RNAs between the two parents, and between the hybrids and parents.**

**(A)** and **(B)** Two hypothetical models of new CHH DMRs induced in WTF1. **(C)** 24-nt small interfering RNA (siRNAs) of new CHH DMRs in Model A. **(D)** 24-nt siRNAs of new CHH DMRs in Model B. **(E)** Ratios of 24 nt siRNAs of high parent to low parent, and of high-parent allele to low-parent allele at the new CHH DMRs in Model A. **(F)** Ratios of 24-nt siRNAs of high parent to low parent, and of high-parent allele to low-parent allele at the new CHH DMRs in Model B. HP, high parent (parent with higher methylation). LP, low parent (parent with lower methylation). MPV, the middle parent value. **, $P < 0.01$, Student's $t$ test. DMRs, differentially methylated regions.

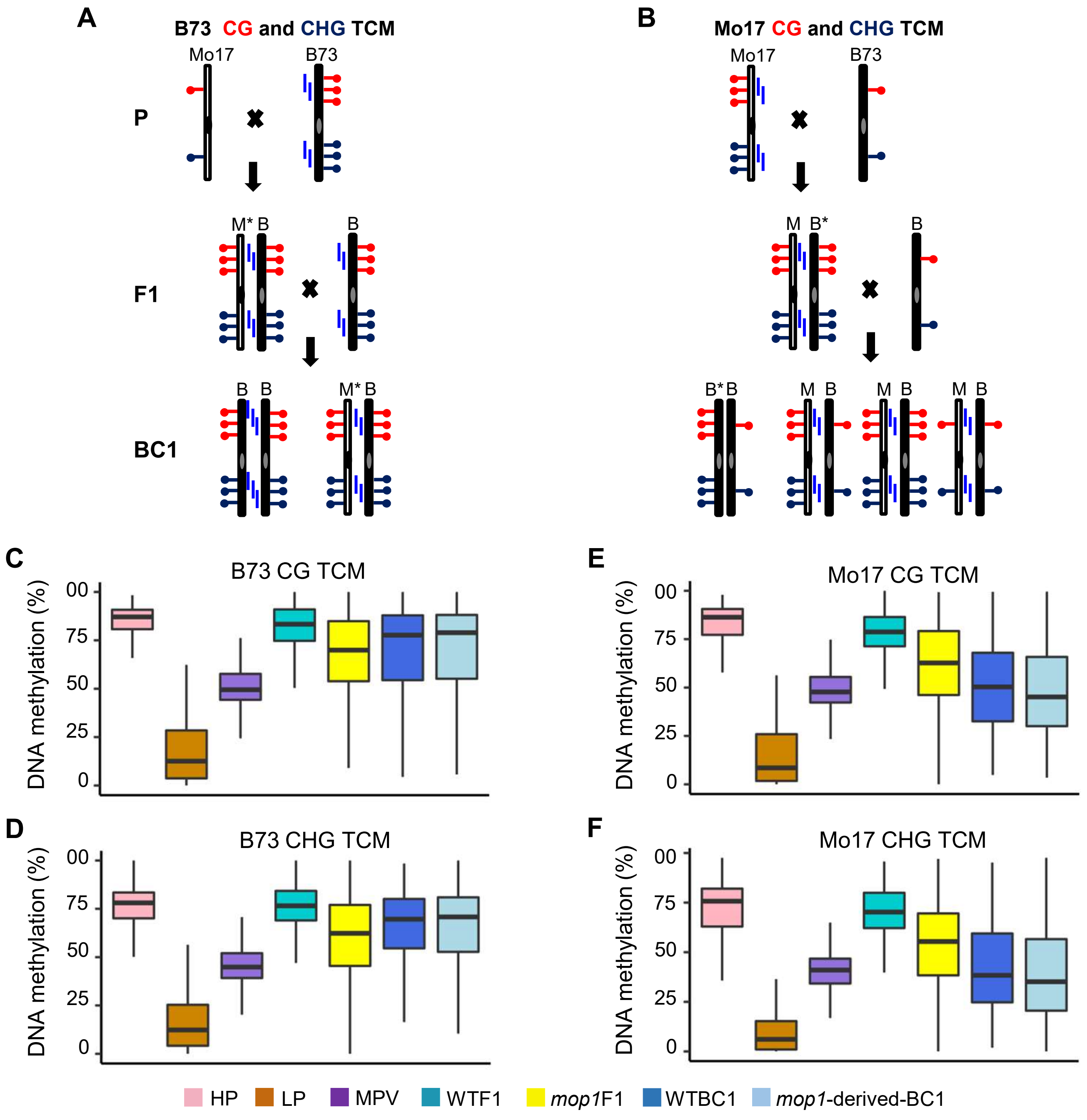


**S15 Fig. Inheritance of newly triggered methylation at CG and CHG TCM DMRs in the backcrossed generation**.

**(A)** Hypothetical model of maintenance of B73 CG and CHG TCM DMRs. Asterisk denotes the newly converted (methylated) allele. **(B)** Hypothetical model of maintenance of Mo17 CG and CHG TCM DMRs. **(C)** Methylation changes of B73 CG TCM DMRs. **(D)** Methylation changes of B73 CHG TCM DMRs. **(E)** Methylation changes of Mo17 CG TCM DMRs. **(F)** Methylation changes of Mo17 CHG TCM DMRs. DMRs, differentially methylated regions. TCM, *trans*-chromosomal methylation. Homo, homozygous. Hetero, heterozygous. WTBC1, Mo17/B73;+/+ × B73. *mop1*-derived BC1, Mo17/B73;*mop1/mop1* × B73.

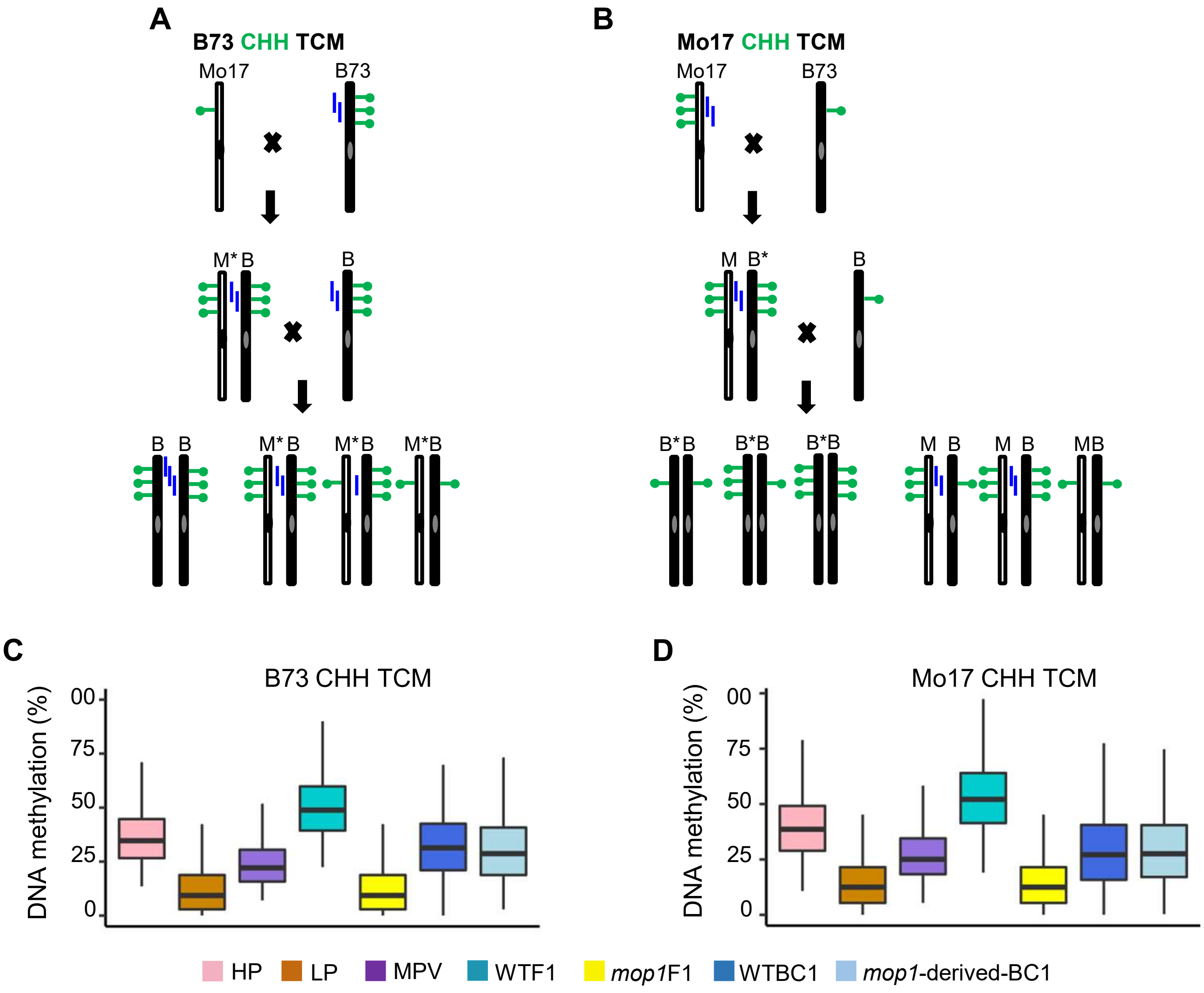


**S16 Fig. Inheritance of newly triggered methylation at CHH TCM DMRs in the backcrossed generation**.

**(A)** Hypothetical model of maintenance of B73 CHH TCM DMRs. Asterisk denotes the newly converted (methylated) allele. **(B)** Hypothetical model of maintenance of Mo17 CHH TCM DMRs. **(C)** Methylation changes of B73 CHH TCM DMRs. **(D)** Methylation changes of Mo17 CHH TCM DMRs. DMRs, differentially methylated regions. TCM, *trans*-chromosomal methylation. Homo, homozygous. Hetero, heterozygous. WTBC1, Mo17/B73;+/+ × B73. *mop1*-derived BC1, Mo17/B73;*mop1/mop1* × B73.

**S1 Table. The summary of raw reads of different samples.**

| Sample name | Smaple ID | Genotype | WGBS | Small RNAs | RNA-seq |
|---|---|---|---|---|---|
| B73 | 1898-12 | B73, parent | 135,703,170 | 39,491,695 | 43,390,794 |
| | 1898-24 | B73, parent | 138,962,192 | 28,301,778 | 40,410,190 |
| Mo17 | 1932-9 | Mo17, parent | 133,751,646 | 32,221,373 | 45,031,488 |
| | 1932-11 | Mo17, parent | 138,933,460 | 24,678,975 | 42,558,268 |
| WTF1-*Mop1* | GP1-40 | Mo17/B73;+/+, wild type F1 hybrid | 223,549,992 | 26,031,157 | 66,825,672 |
| | GP5-19 | Mo17/B73;+/+, wild type F1 hybrid | 220,604,148 | 27,963,678 | 66,902,702 |
| *mop1* F1 | GP1-5 | Mo17/B73;*mop1-1* /*mop1-1* , *mop1* mutant F1 hybrid | 220,604,148 | 25,261,773 | 67,497,392 |
| | GP5-40 | Mo17/B73;*mop1-1* /*mop1-1* , *mop1* mutant F1 hybrid | 182,065,938 | 27,735,347 | 66,902,702 |
| WTBC1 | 2146-8 | wild type BC1 | 54,092,722 | | |
| | 2146-24 | wild type BC1 | 53,083,340 | | |
| | 2146-30 | wild type BC1 | 46,123,474 | | |
| | 2146-49 | wild type BC1 | 44,990,674 | | |
| | 2148-16 | wild type BC1 | 46,090,340 | | |
| | 2148-20 | wild type BC1 | 52,256,536 | | |
| | 2148-22 | wild type BC1 | 43,342,938 | | |
| | 2148-31 | wild type BC1 | 53,767,458 | | |
| *mop1* -derived-BC1 | 2145-21 | *mop1/+* BC1 | 48,516,778 | | |
| | 2145-22 | *mop1/+* BC1 | 57,738,922 | | |
| | 2145-29 | *mop1/+* BC1 | 47,834,240 | | |
| | 2145-33 | *mop1/+* BC1 | 54,457,578 | | |
| | 2147-30 | *mop1/+* BC1 | 52,242,914 | | |
| | 2147-32 | *mop1/+* BC1 | 46,433,494 | | |
| | 2147-3 | *mop1/+* BC1 | 45,369,542 | | |
| | 2147-33 | *mop1/+* BC1 | 56,508,958 | | |

**S2 Table. The overall patterns of cytosine methylation in parents, WTF1, and mutant F1.**

| Features | B73 | | Mo17 | | WTF1 | | *mop1* F1 | |
|---|---|---|---|---|---|---|---|---|
| | rep1 | rep2 | rep1 | rep2 | rep1 | rep2 | rep1 | rep2 |
| Mapping efficiency (%) | 73.4 | 73.7 | 53.6 | 54 | 58.8 | 59.7 | 59.9 | 58.3 |
| Total C (%) | 25.6 | 24.6 | 25.2 | 24.7 | 30.6 | 29.7 | 28.1 | 30.9 |
| CG (%) | 87.4 | 86.3 | 84.2 | 85.6 | 85.5 | 81.5 | 82.3 | 84.3 |
| CHG (%) | 71.1 | 69.7 | 68.1 | 69.2 | 73.3 | 69 | 68.9 | 72.5 |
| CHH (%) | 1.5 | 1.5 | 1.5 | 1.6 | 2.4 | 2 | 1.8 | 2 |

**S3 Table. DMRs identified between parents (B73 and Mo17).**

| Parental DMRs | Number of DMRs | DMR length (bp) | Number of B73 hyper DMRs | Number of Mo17 hyper DMRs |
|---|---|---|---|---|
| CG | 7107 | 371.2 | 5602 | 1505 |
| CHG | 9045 | 478.9 | 6099 | 2946 |
| CHH | 13307 | 74.8 | 3021 | 10286 |

**S4 Table. Number of changed DMRs in the other two cytosine contexts at the *mop1* -affected CG, CHG, and CHH DMRs.**

| *mop1* affected DMRs | No. of changed DMRs in the other two cytosine contexts |
|---|---|
| CG DMRs (N = 118) | 37 (CHG) |
| | 3 (CHH) |
| CHG DMRs (N = 153) | 32 (CG) |
| | 9 (CHH) |
| CHH DMRs (N = 1048) | 72 (CG) |
| | 181 (CHG) |

**S5 Table. Differentially expressed genes involved in the transcriptional gene silencing pathway between parents.**

| Gene ID | B73 (rep1) | B73 (rep2) | Mo17 (rep1) | Mo17 (rep2) | $\log_2$ (Mo17/B73) | Gene name |
|---|---|---|---|---|---|---|
| Zm00001d003378 | 1061.20 | 1193.00 | 1645.96 | 2105.39 | 0.74 | RDR2 |
| Zm00001d008249 | 2123.36 | 2735.74 | 3614.57 | 4069.70 | 0.66 | AGO4 |
| Zm00001d010646 | 962.28 | 912.59 | 2126.88 | 2105.39 | 1.17 | SUVH2 |
| Zm00001d019905 | 1235.03 | 1152.21 | 2084.00 | 2089.71 | 0.81 | AGO6 |
| Zm00001d024677 | 119.08 | 172.32 | 453.35 | 412.46 | 1.57 | DRD1 |
| Zm00001d038113 | 181.51 | 171.30 | 358.39 | 249.82 | 0.79 | CLSY3 |
| Zm00001d048516 | 281.39 | 216.17 | 412.51 | 424.21 | 0.75 | DRM2 |
| Zm00001d049884 | 826.87 | 755.57 | 1478.50 | 1321.62 | 0.82 | SUVR2 |

**S6 Table. Differentially expressed genes involved in the transcriptional gene silencing pathway between MPV and F1.**

| Gene ID | MPV rep1 | MPV rep2 | WTF1 rep1 | WTF1 rep2 | $log_2$ (WTF1/MPV) | Gene name |
|---|---|---|---|---|---|---|
| Zm00001d010646 | 1475.41 | 1489.65 | 3050.98 | 2723.15 | 0.96 | SUVH2 |
| Zm00001d010719 | 1533.78 | 1360.45 | 5122.36 | 5529.26 | 1.88 | KTF1 |
| Zm00001d024677 | 271.74 | 288.73 | 568.65 | 604.00 | 1.07 | DRD1 |
| Zm00001d026421 | 419.09 | 415.97 | 837.05 | 910.46 | 1.07 | NRPD2/NRPE2 |
| Zm00001d038644 | 1039.11 | 939.59 | 5681.91 | 4329.48 | 2.34 | KTF1 |
| Zm00001d053872 | 2541.31 | 2379.33 | 6314.25 | 6057.16 | 1.33 | NRPE1 |

**S7 Table. Inheritance of CG and CHG TCM and TCdM in the backcrossed generation (BC1).**

| DMRs | Total number | Methylation difference between WTF1 and WTBC1 less than 10% | Percentage | Average |
|---|---|---|---|---|
| B73 CG TCdM | 339 | 99 | 29.20% | 35.14% |
| Mo17 CG TCdM | 112 | 46 | 41.07% | |
| B73 CG TCM | 904 | 316 | 34.96% | 24.66% |
| Mo17 CG TCM | 188 | 27 | 14.36% | |
| B73 CHG TCdM | 235 | 74 | 31.49% | 43.98% |
| Mo17 CHG TCdM | 147 | 83 | 56.46% | |
| B73 CHG TCM | 872 | 294 | 33.72% | 25.98% |
| Mo17 CHG TCM | 329 | 60 | 18.24% | |

**S8 Table. Inheritance of CHH TCM and TCdM in the backcrossed generation (BC1).**

| DMRs | Total number | Methylation difference between WTF1 and WTBC1 less than 5% | Percentage | Average |
|---|---|---|---|---|
| B73 CHH TCdM | 64 | 22 | 34.38 | 37.66 |
| Mo17 CHH TCdM | 237 | 97 | 40.93 | |
| B73 CHH TCM | 197 | 26 | 13.20 | 10.52 |
| Mo17 CHH TCM | 714 | 56 | 7.84 | |
